# Differential submergence tolerance between juvenile and adult Arabidopsis plants involves the ANAC017 transcription factor

**DOI:** 10.1101/2020.02.12.945923

**Authors:** Liem T. Bui, Vinay Shukla, Federico M. Giorgi, Alice Trivellini, Pierdomenico Perata, Francesco Licausi, Beatrice Giuntoli

## Abstract

Plants need to attune stress responses to the ongoing developmental programs to maximize their efficacy. For instance, successful submergence adaptation is often associated to a delicate poise between saving resources and their expenditure to activate measures that allow stress avoidance or attenuation. We observed a significant decrease in submergence tolerance associated with aging in *Arabidopsis thaliana*, with a critical step between two and three weeks of post-germination development. This sensitization to flooding was concomitant with the transition from juvenility to adulthood. Transcriptomic analyses indicated that a group of genes related to ABA and oxidative stress response was more expressed in juvenile plants than in adult ones. These genes are induced by endomembrane tethered ANAC factors that were in turn activated by submergence-associated oxidative stress. A combination of molecular, biochemical and genetic analyses showed that these genes are located in genomic regions that move towards a heterochromatic state with adulthood, as marked by lysine 4 dimethylation of histone H3. We concluded that, while the mechanism of flooding stress perception and signal transduction were unaltered between juvenile and adult phases, the sensitivity that these mechanisms set into action is integrated, via epigenetic regulation, into the developmental programme of the plant.

## Introduction

Plants undergo transitory developmental phases before engaging in reproduction. Such intermediate steps are required to establish offspring production when metabolic resources are amply available, and ensure seed dispersal under convenient environmental conditions. Typically, vegetative development towards full sexual maturation progresses through discrete juvenile and adult stages (Poethig, 2013). The main difference between the two is the competence to integrate endogenous and environmental cues to initiate the next developmental transition: flowering. Often, these phases are marked by distinctive anatomical traits. This can be expressed as extremely divergent dimorphism, originally termed heteroblasty, as in some perennial woody species (Hildebrand, 1875; James & Bell, 2001). In the small annual plant *Arabidopsis thaliana*, the transition between juvenility and adulthood entails the formation of abaxial foliar trichomes, leaf elongation and serration and decrease in newly produced cells (Telfer *et al.*, 1997; Tsukaya *et al.*, 2000).

SQUAMOSA PROMOTER BINDING LIKE PROTEINS (SBP/SPLs) act as conserved integrators between environmental cues, such as photoperiod, and the metabolic state of the plant to drive these morphological changes (Huijser & Schmid, 2011). Active sugar metabolism and carbon availability have been shown to control SPL abundance both at the transcriptional and posttranscriptional level (Matsoukas *et al.*, 2013; Yu *et al.*, 2013). A number of proteins involved in DNA methylation and chromatin remodelling mediate the former layer of regulation, whereas the second is achieved via the microRNAs miR156 and miR157 that cleave and prevent translation of SPL transcripts (Wingler, 2018).

Beyond acting as exogenous cues that promote stage-specific morphological alterations and acquisition of reproductive competence, environmental stresses also exert variable effects on plant life depending on the developmental stage at which they are experienced. This can be explained by the intrinsic tolerance of organs produced at different developmental stages, as well as changes in the response elicited by the perception of the stimulus, as observed in the case of drought, heat and cold stress (Lim *et al.*, 2014; Marias *et al.*, 2017; Kanojia & Dijkwel, 2018). Understanding stress tolerance mechanisms is extremely relevant to minimize crop yield reduction by adverse environmental conditions. Indeed, several mechanisms that improve plant tolerance and fitness have been identified and are currently exploited (Bechtold & Field, 2018).

Despite the relevant threat to worldwide agricultural production posed by flooding-related events (Voesenek & Bailey-Serres, 2015), the biochemical and molecular bases for tolerance to flooding are less well established when compared to the abiotic stress conditions mentioned above. This limitation is connected to the complex nature of the flooding stress by itself. First, submergence results in the concomitant depletion of oxygen available for respiration, accumulation of carbon dioxide, ethylene and hydrogen sulphide, and enhanced mobilisation of reduced phytotoxic compounds (Bailey-Serres & Colmer, 2014). Moreover, once the water level recedes, de-submerged plants are exposed to a dehydration-like stress that jeopardizes their fitness, if not their very survival (Yeung *et al.*, 2019). Flooding tolerance mainly relies on two alternative strategies. The first enables stress avoidance by investing the available resources in rapid elongation and leaf petiole reorientation to emerge from the water surface. The second, instead, reduced ATP and carbohydrates consumption restraining metabolism to the essential reactions during the stress, to resume active growth and support reproduction in the aftermath (Loreti *et al.*, 2016). The alternative activation of either strategy is genetically determined and entails the perception and integration of several cues.

In higher plants, cellular oxygen levels are monitored by the enzymatic class of N-terminal cysteine oxidases of plants (PCOs) (Weits *et al.*, 2014; White *et al.*, 2017), which control the activity of the group VII Ethylene Responsive Factors (ERF-VII) (Gibbs *et al.*, 2011; Licausi *et al.*, 2011), key regulators of the anaerobic metabolism and oxidative stress response (Giuntoli & Perata, 2017). The stability of these transcriptional regulators is directly linked to oxygen availability, since under aerobic conditions oxidation of an exposed cysteine at their N-terminus by PCO directs them towards degradation via the N-degron pathway (van Dongen & Licausi, 2015). Besides the ERF-VIIs, the accumulation of reactive oxygen and nitrogen species (ROS and RNS) also leads to the activation of additional transcription factors that control the homeostasis of these potentially harmful molecular species, including members of the Heat Shock Factor (HSFs) and NAC families (Gonzali *et al.*, 2015). Recently, this latter family of transcriptional regulators has attracted the attention of the flooding community. A particular subgroup of endoplasmic reticulum (ER)-bound NAC factors can be activated by ROS via endoproteolytic cleavage, whereby the release of N-terminal TF fragments into the nucleus activates genes involved in ROS homeostasis and mitochondrial metabolism (De Clercq *et al.*, 2013; Ng *et al.*, 2013; Meng *et al.*, 2019).

In the present study, we investigate the dependence of Arabidopsis tolerance to flooding conditions on plant age. Having characterized a differential stress response connected with developmental transitions, we propose an ER-tethered NAC TF as a key regulator the molecular mechanisms underneath the phenomenon.

## Materials and methods

### Plant material and growth conditions

*Arabidopsis thaliana* Col-0 ecotype was used as wild type in all the experiments. The T-DNA insertional mutants *anac017* (SALK_022174) and *abi2-1* (NASC ID N23), and a *35S:miR156* over-expressing line (N9952; (Wang *et al.*, 2009)) were obtained from the European Arabidopsis Stock Centre (NASC). Homozygous *anac017* plants were confirmed by standard PCR, using the gene-specific primers insANAC017_F (5’-GGGCTCCTAGTGGTGAGCGGACTGA) and insANAC017_R (5’-CTCATCGATATCCTCTAACTGAAGA), and LBb1 (5’-ATTTTGCCGATTTCGGAAC) as a T-DNA specific primer. Plants were germinated and grown on peat (Hawita Flor), using a peat:sand ratio 3:1, after stratification at 4°C in the dark for 48 h. Plants were maintained in growth chamber under 23°C/18°C (day/night) temperature cycle, 80 µmol photon m^-2^ s^-1^ light intensity and neutral photoperiod (12 h/12 h light/dark). For axenic growth conditions, seeds were surface sterilized and germinated on half strength Murashige and Skoog (Duchefa) basal medium (pH 5.7) supplemented with 5 g l^-1^ sucrose and 8 g l^-1^ plant agar (Duchefa). All experiments entailing the comparison between juvenile and adult plants referred to individuals at the developmental stage defined, respectively, as 1.06 and 1.12 by (Boyes, 2001), corresponding to two and three weeks of growth in our experimental conditions.

### Submergence experiments and chemical treatments

At end of the light phase, soil-grown plants were submerged with tap water in glass tanks, until the water surface reached approximately 20 cm above the rosettes; water was equilibrated to room temperature along the previous day. Flooding was protracted in the absence of illumination for the specified duration, while control plants were maintained in darkness in air. After submergence, plants were either sampled for the specific subsequent analyses or, moved back to normal photoperiodic conditions and allowed to recover for one week before survival scoring. Plants that were able to produce new leaves in the recovery period were categorized as alive, on the opposite they were scored as dead. The number of plants deployed in each submergence experiment is indicated in the corresponding sections, with a minimum of four replications with 15 plants each. Chemical treatments were administered by spraying with 100 nM antimycin A in 0.001% ethanol 6 h before submergence, or 10 µM ABA in 0.1% ethanol 3 h in advance, whereas control plants were sprayed with the respective solvent.

### Microarray analysis

Total RNA was extracted using the Spectrum™ Plant Total RNA Kit (Sigma-Aldrich) from a pool of three Col-0 plants at the juvenile or adult stage. Two biological replicates were used. Hybridization against the Arabidopsis Gene 1.0 ST Array, washing, staining and scanning were performed by Atlas Biolabs GmbH. Normalization of the raw microarray data and extraction of signal intensities were carried out through the Robust MultiArray methodology (Giorgi *et al.*, 2010). An empirical Bayes method to shrink the probe-wise sample variances was applied to the data, and then differential expression analysis was carried out using the Limma R package (Ritchie *et al.*, 2015). Microarray data were deposited in the Gene Expression Omnibus repository with the accession number GSE137866.

### RNA extraction and real time qPCR analyses

Equal amounts of total RNA, extracted as above, were subjected to DNase treatment, carried out using the RQ1-DNase kit (Promega), and were reverse transcribed into cDNA using the iScript™ cDNA Synthesis Kit (Biorad). was performed on 15 ng cDNA were used for real time qPCR amplification with the ABI Prism 7300 sequence detection system (Applied Biosystems), using iQSYBR Green Supermix (Biorad). All primers are listed in Table S1. Relative gene expression was calculated using the 2^-ΔΔCt^ Method (Livak & Schmittgen, 2001), based on the housekeeping gene *Ubiquitin10* (*At4g05320*).

### Production of *RFP-ANAC017-GFP* transgenic lines in *A. thaliana*

In order to obtain an RFP-ANAC017-GFP fusion construct, the full length coding sequence of *ANAC017* (1671 bp, devoid of the stop codon) was amplified from an Arabidopsis cDNA sample with the primers gwANAC017_F (5’-CACCATGGCGGATTCTTCACCCGA) and ANAC017-XhoI-HindIII_R (5’-AAGCTTAGAGCTCGAGGTCTTTCAAGAGAAGA). The template was obtained from shoot material extracted as described above and processed into cDNA by use of RQ1-DNase (Promega) and SuperScript III Reverse Trascriptase kit (Thermo-Fisher Scientific). The sequence was subsequently cloned in the pENTR/D-TOPO vector (Thermo-Fisher Scientific), to generate a pENTR-ANAC017 plasmid. The GFP coding sequence was amplified from the template plasmid pK7FWG2 (Karimi *et al.*, 2002), with the primers XhoI-eGFP_F (5’-CACCGCGCTCGAGATGGTGAGCAAGGGCGAGG) and eGFP-HindIII_R (5’-CGCAAGCTTTTACTTGTACAGCTCG), and cloned into pENTR to obtain a pENTR-GFP plasmid. Subsequently, the two coding sequences were fused in frame into a new pENTR-ANAC017-GFP vector, upon digestion of the two starting entry plasmids with XhoI and HindIII, and ligation of the resulting pENTR-ANAC017 linearized vector with the GFP fragment. Finally, the Gateway LR clonase II enzyme mix (Thermo-Fisher Scientific) was used to recombine pENTR-ANAC017-GFP with the N-terminal tagging vector pUBN-RFP (Grefen *et al.*, 2010). The resulting plant expression vector pUBQ10:RFP-ANAC017-GFP was employed for Arabidopsis stable transformation, achieved through the floral dip method (Zhang *et al.*, 2006). First generation transformants (T_1_) were selected upon germination on MS medium plates containing the herbicide glufosinate-ammonium (PESTANAL**®**, Sigma-Aldrich). Transgene expression was assessed by real time qPCR using the primers ANAC017_F and ANAC017_R listed in Table S1. Single-copy transgene insertion was verified by segregation of the T_2_ progeny on herbicide-containing plates. Homozygous T_3_ or next generation plants were used for the following experiments.

### Western blot

Total protein from shoot tissues of *pUBQ10:RFP-ANAC017-GFP* plants was extracted in a buffer containing 50 mM Tris-HCl pH 7.6, 1 mM EDTA, 100 mM NaCl, 2% SDS and 0.05% Tween-20. Fifty µg proteins from the extracts, quantified with the Pierce BCA Protein Assay Kit (Thermo-Fisher Scientific), were separated by SDS–PAGE on 10% polyacrylamide Bis-Tris NuPAGE midigels (Thermo-Fisher Scientific) and transferred onto a polyvinylidene difluoride membrane by means of the Trans-Blot Turbo System (Bio-Rad). A monoclonal anti-GFP antibody (Roche, cat. no. 11814460001) was used at 1:3000 dilution and combined with a 1:20000 rabbit anti-mouse secondary antibody (Agrisera, cat. no. AS09627) to detect the Δ533ANAC017:GFP protein fragment. A rat monoclonal anti-RFP antibody (Chromotek, cat. no. 5F8) was used at 1:1000 dilution and combined with a 1:5000 donkey anti-rat secondary antibody (Agrisera, cat. no. AS10947) to detect the RFP:ANAC017Δ24 protein fragment. All antibodies were diluted in 4% milk in TBS-T. Blots were incubated with the LiteAblot Turbo Extra Sensitive Chemiluminescent Substrate (EuroClone) and imaged with UVP VisionWorks LS (Ultra-Violet Products). Equal loading of total protein samples was checked by Amido-black staining of the membrane (Mithran *et al.*, 2014).

### ROS staining and quantification

Reactive oxygen species (ROS) production was visualized in juvenile and adult plants treated in dark submergence conditions for 24 h, starting at the end of the light phase, or sampled before the onset of the treatment. ROS staining and detection was carried out using methods described by Daudi and O’Brien (Daudi & O’Brien, 2012). Briefly, collected plants were stained in freshly prepared 1 mg ml^-1^ 3,3’-diaminobenzidine solution (DAB, Sigma-Aldrich) neutralized in 10 mM Na_2_HPO_4_ for 12 h, then chlorophyll was bleached in an ethanol:acetic acid:glycerol=3:1:1 solution with boiling (95°C for 15 min). After that, plants were imaged by means of an optical scanner and the acquired images were processed with Image J (Rueden *et al.*, 2017) to obtain a digital quantification of pixel intensity relative to the background and normalize it to the area of the object.

### ABA quantification

Extraction and determination of abscisic acid content in Arabidopsis shoots were performed as described in (Woo *et al.*, 2011). Briefly, 200–300 mg plant material was frozen in liquid nitrogen and ground using a Mixer-Mill (Qiagen). The homogenate was extracted with 1–2 ml ABA extraction buffer (10 mM HCl and 1% w/v polyvinyl polypyrrolidone in methanol), with continuous mixing overnight at 4 °C. After centrifugation, the supernatant was neutralized with 1 M NaOH, and ABA levels were quantified in the extract using the Phytodetek-ABA kit (AGDIA Inc.), following the manufacturer’s protocol. Raw values for ABA levels were standardized on sample fresh weight and extraction volume.

### Genomic DNA methylation analysis

Genomic DNA was extracted from 40 mg leaf material of juvenile or adult *A. thaliana* Col-0 plants kept under control growth conditions, using the Wizard Genomic DNA Purification Kit (Promega). Cytosine methylation on target loci was assessed by Methylation-Sensitive Restriction Enzyme qPCR (MSRE-qPCR) (Hashimoto *et al.*, 2007) using the methylation sensitive endonucleases HpaII and SalI (Thermo Fisher Scientific). Full information related to target loci identity and location of the target CpG sites evaluated is reported in Table S2. Eight hundred ng DNA was digested using 1 µl of the respective restriction enzyme (10 U HpaII, or 40 U SalI), or an equal volume of 50% glycerol for mock digestions, in 20 µl reaction volume containing the appropriate buffer according to the manufacturer’s recommendations. Reactions were incubated over night at 37°C, followed by enzyme inactivation at 65°C for 30 min. Enzyme-treated and mock-treated samples were diluted down to 10 ng µl^-1^ and 1 µl DNA was amplified by qPCR using 200 nM of each primer and Power Up SYBR Green Master Mix (Thermo Fisher Scientific). Details on the primers used for the analysis can be found in Table S2. Amplifications were carried out with an ABI Prism 7300 sequence detection system (Applied Biosystems) with a standard cycling protocol. The methylation level in each sample is expressed as relative amplification level (cut/uncut ratio) between enzyme- and mock-treated DNA, calculated as 2^-ΔCt^ values, where ΔC_t_=C_t_(cut)-C_t_(uncut).

### Chromatin immuno-precipitation

The extent of histone methylation in juvenile or adult leaf samples was assessed in Col-0 plants by chromatin immuno-precipitation with rabbit polyclonal Anti-Histone H3 (di methyl K4) (Abcam, cat. no. ab7766) or Anti-Histone H3 (tri methyl K27) antibody (Abcam, cat. no. ab195477) (5 µg specific antibody added to each sample). Five biological replicates were used, each obtained from 500 mg fresh tissue. The ChIP assay was performed according to the protocol described in Giuntoli et al. (2017). The primers used to specifically quantify genomic DNA abundance by qPCR in immune-purified and input samples are listed in Table S3.

### Statistical analyses

All ANOVA and Kaplan-Meier survival analyses were carried out with the aid of the GraphPad Prism version 6.01 for Windows (GraphPad Software, La Jolla California USA, www.graphpad.com). Additional details related to individual experiments are provided in the corresponding sections and legends.

## Results

### Arabidopsis tolerance to submergence conditions is age-dependent

We systematically assessed the flooding tolerance of Arabidopsis plants (Columbia-0 ecotype) at different ages. Two, three, four or five weeks after germination, plants corresponding to stage 1.06, 1.12, 3.70 or 3.90 of vegetative development (Boyes et al. 2001) were subjected to submergence for two to four days, with incremental steps of 12 h. Survival was scored a week after de-submergence. Two week-old plants clearly exhibited superior performance under prolonged submergence (Fig. 1a) that was further confirmed by their higher survival rates as compared to older plants (Fig. 1b).

**Fig. 1.**
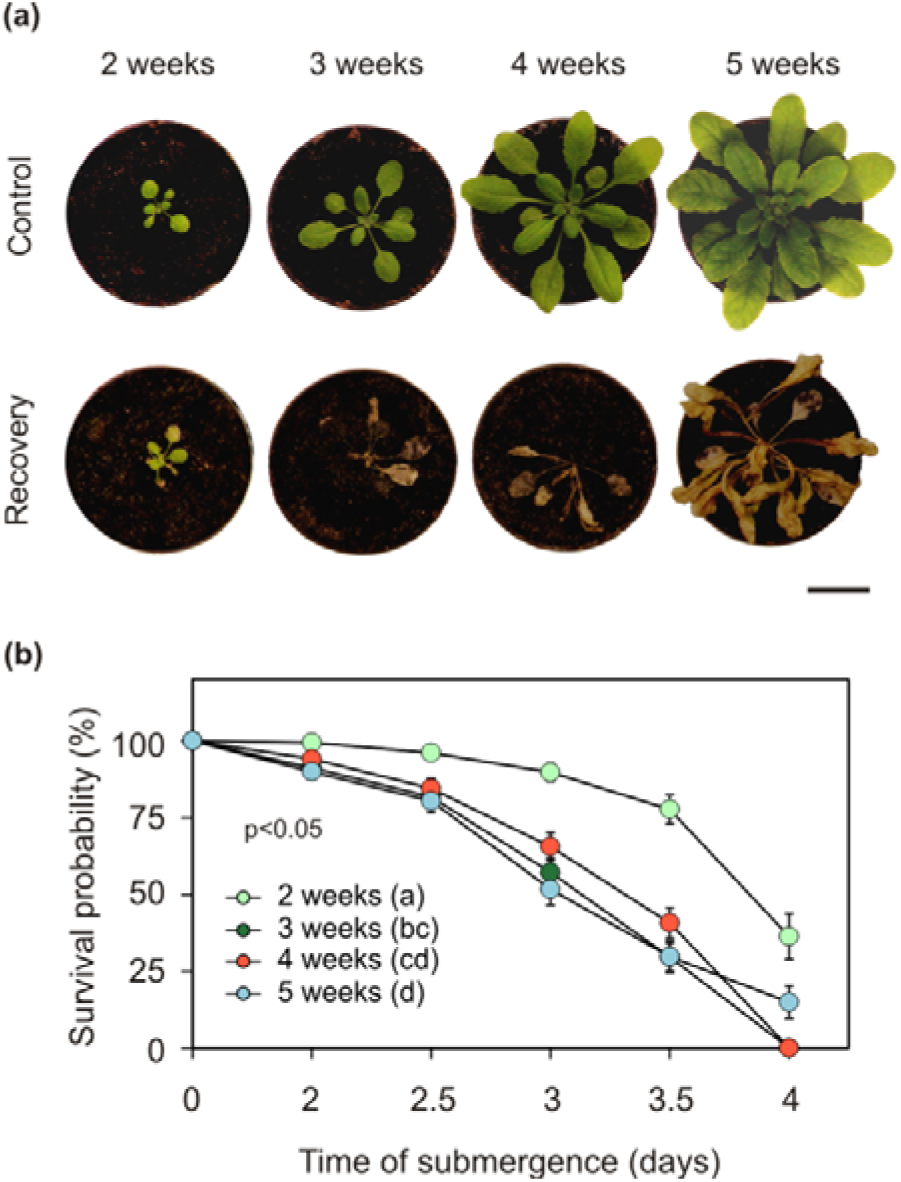
Submergence tolerance at different developmental stages of Arabidopsis rosettes. **(a)** Plant phenotypes immediately before the flooding treatment (“Control”) and one week after recovery following 4 days full submergence (“Recovery”). Scale bar, 2 cm. **(b)** Survival probability of Arabidopsis plants at four different ages when subjected to flooding (n=25 for each combination of plant age and stress duration). Data are mean ± S.E. from four independent experiments. Letters in brackets indicate statistically significant difference among the survival curves, according to the Kaplan-Meier test (p<0.05).

In the attempt to reduce the space required for the subsequent experiments, we then tested plants grown in pools of equally spaced individuals, in 7 cm pots (Fig. 2a), instead of singularly. In these new conditions, two week-old plants again displayed better flooding tolerance than three week-old ones (Fig. 2a and 2b), demonstrating the suitability of the set-up. In the older plants, the first symptoms of sufferance, such as vitrescence and collapse of leaf tissues, could already be observed at de-submergence (Fig. 2a), although loss of survival capacity, in terms of maintenance of leaf production at the shoot apical meristem, was evident only one week later.

**Fig. 2.**
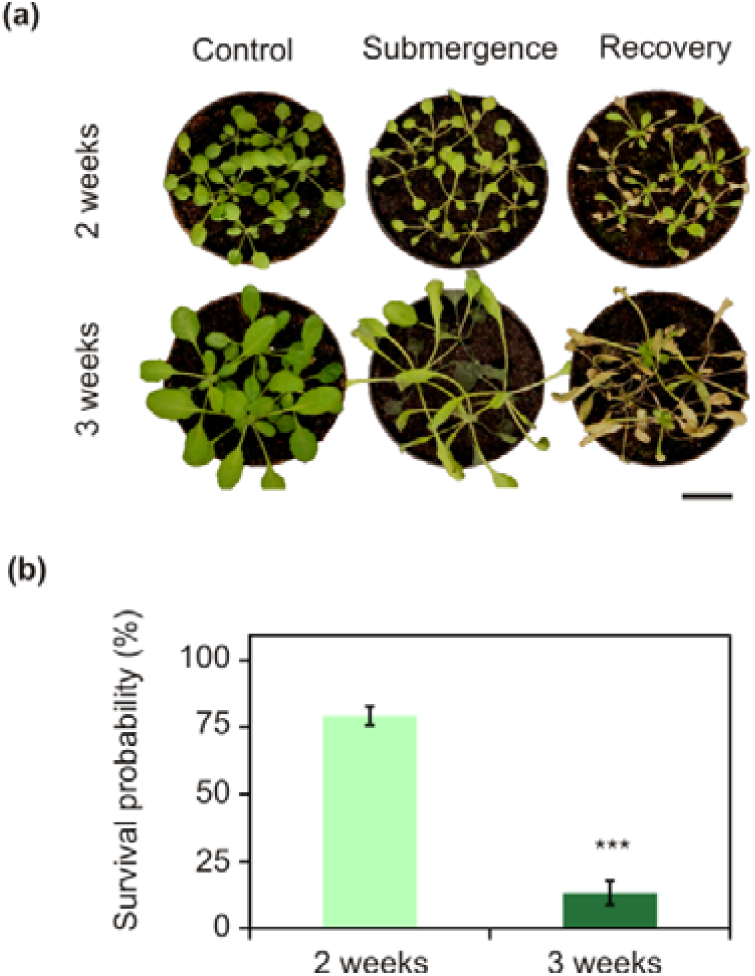
Submergence tolerance of two and three-week-old seedlings grown collectively in pots. **(a)** Phenotypes of two and three week old plants before submergence (left), immediately after being de-submerged at the end of a 72 h-long treatment (centre), and one week after (right). Scale bar, 2 cm. **(b)** Survival rate scored after one week following 72 h submergence (n=30 for 2 weeks of age, n=15 for 3 weeks). Data are present as mean ± S.E (n=4) and asterisks show significant difference according to the Kaplan-Meier test (p<0.01).

Attracted by this phenomenon, we decided to investigate its molecular determinants. Our first observation was that the difference between the two phenological stages encompassed the transition from juvenilily to adulthood. On average, the rosette of two week-old plants consisted of six leaves; one week later, this number doubled (Fig. 3a). In this time interval, moreover, canonical markers of adulthood were displayed, such as the appearance of trichomes in the abaxial side of the eighth leaf and those subsequently produced, and elongation of leaf shape with serrated margins (Fig. 3a).

**Fig. 3.**
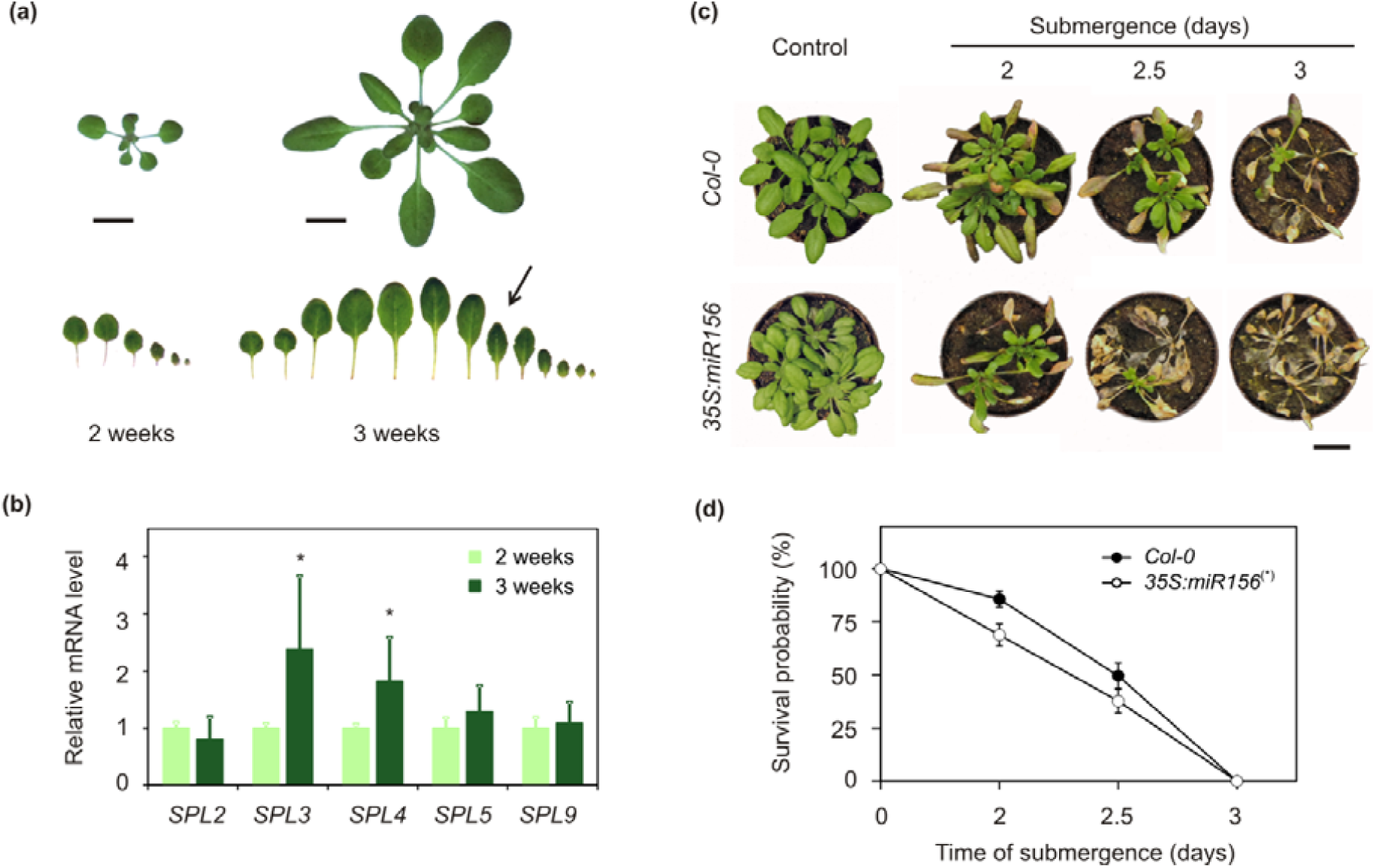
Juvenile to adult phase transition after the second week of growth of Arabidopsis rosettes. **(a)** Size and leaf number of two- and three-week-old plants. True leaves are displayed progressively according to their age, from left to right. The arrow indicates the first leaf with abaxial trichomes, taken as a marker of adulthood. Scale bars, 1 cm. **(b)** Relative mRNA level of *SPL2, 3, 4, 5* and *9* genes in two- and three-week-old plants as assessed by real time qPCR. Data are mean ± S.D., asterisks indicate statistically significant difference at p<0.05 after t-test (n=4). Relative expression was set to 1 in two week-old plants. **(c)** Phenotype in three week-old wild type and *35S:miR156* over-expressors before submergence (left), or immediately after 48, 60 or 72 h flooding. Scale bar, 2 cm. **(d)** Submergence tolerance in the wild-type and *35S:miR156*s (n=30 for each combination of plant age and stress duration), assessed as survival probability. Data are presented as mean ± S.E. (n=4). The asterisk indicates a significant difference in survival (p<0.05), according to the Kaplan-Meier test.

SPL transcription factors have been identified as crucial regulators of this developmental transition (Huijser & Schmid, 2011; Zhang *et al.*, 2015). A survey of fourteen Arabidopsis *SPL* genes in the Genevestigator bioinformatics platform (Hruz *et al.*, 2008) revealed that they are globally highly expressed in vegetative tissues, and that *SPL3* and *4* mRNAs slightly increase from young to adult plants, by a 12% or 19% fold change (Fig. S1). We therefore compared the mRNA levels of a subset of *SPL* genes in shoot tissues of two and three-week-old plants by means of realtime qPCR. In our hands, *SPL3* and *SPL4* mRNA levels showed a marked and significant rise in older plants, while *SPL2, 5* and *9* showed as expected no detectable fluctuations (Fig. 3b). Hence, based on the behaviour of the phase transition markers *SPL3* and *4*, here onwards we refer to two week-old plants as juvenile and three week-old plants as adult.

The surge of *SPLs*’ expression that drives the achievement of adulthood is known to be enabled by suppression of miR156 expression, whereas it overexpression delays the dismissal of juvenile traits. We therefore tested if transgenic plants with ectopic constitutive expression of miR156 also retained higher submergence tolerance. Unexpectedly, three week-old *35S:miR156* plants (Wang et al., 2009), characterised by a prolonged juvenile phase, showed higher sensitivity than adult wild type plants of the same age, both visible at the moment of de-submergence (Fig. 3c) and after recovery (Fig. 3d). We concluded that ectopic miR156 expression exerted a detrimental effect that overcame the improved tolerance to submergence associated with juvenility.

### Molecular responses to hypoxia do not differ between juvenile and adult plants

Once discarded a direct involvement of miR156 in the different tolerance of juvenile and adult plants, we turned to evaluate whether its molecular bases could be revealed by the expression of marker genes. We analysed a number of markers of anaerobic responses (*ADH, PDC1, Hb1, SAD6, HRA1*), ROS scavenging (*APX1*), and sugar starvation (*DIN6, ATL8, TPS8, KMD4*) in juvenile and adult plants subjected to 12 and 24 h of flooding. Bearing in mind the relevance of both submergence and post-submergence adaptive responses (Yeung *et al.*, 2019), we also collected samples 12 and 24 h after de-submergence. Juvenile and adult plants did not exhibit differences in the activation of core low oxygen-responsive genes, which in plants of both ages were strongly upregulated during flooding and equally abated after 24 h re-oxygenation (Fig. 4 and Table S4). Starvation-related genes, instead, exhibited interestingly lower expression in juvenile plants after 24 h submergence as compared with adult ones, but not at the earlier time point (Fig. 4 and Table S4).

**Fig. 4.**
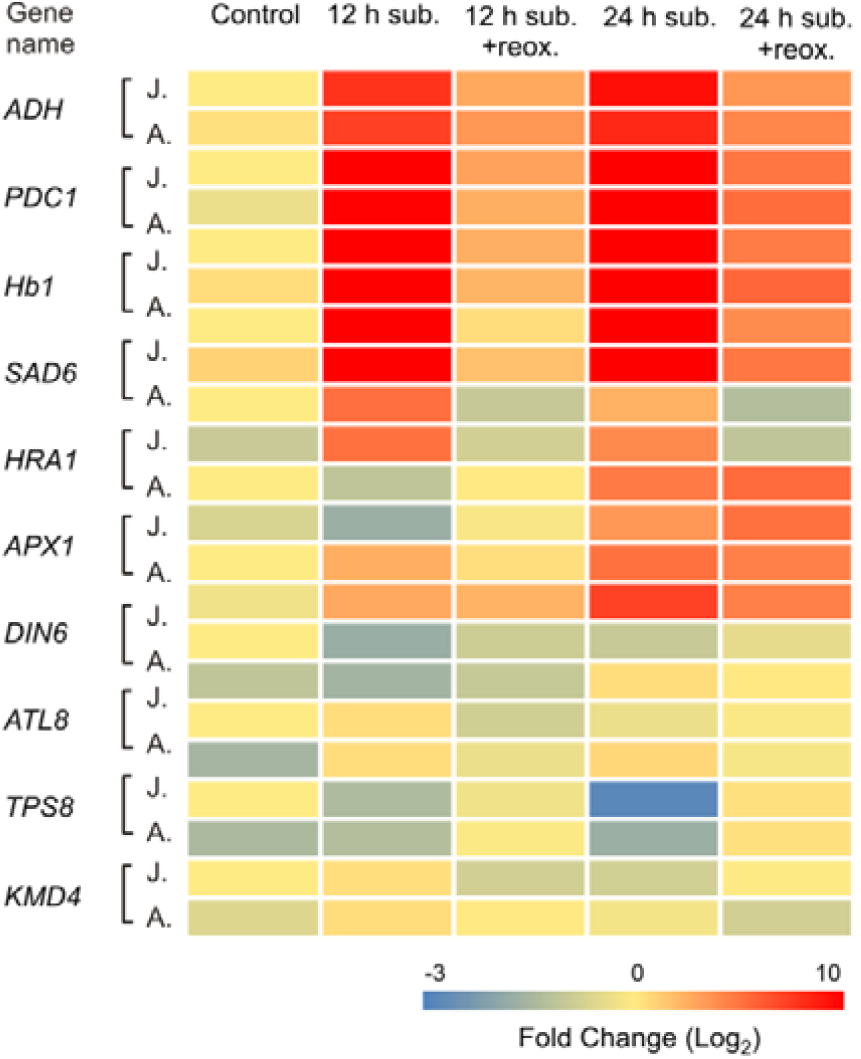
Expression of marker genes for submergence-related processes in treated juvenile or adult plants. Heatmap representation of relative mRNA levels, assessed by real time qPCR, in juvenile (“J.”) or adult plants (“A.”). Transcript abundance was measured after 12 h or 24 h dark submergence, or after 24 h recovery in fully aerated conditions (“+reox.”) following either treatment. Data are mean ± S.D. (n=4) and are expressed, for each age, as fold change (Log_2_ transformed) to the expression in juvenile samples taken immediately before the start of submergence. Supporting values can be found in Table S4. All primers used are listed in Table S1.

### Transcriptome-wide comparison of juvenile and adult plant responses to submergence

Since the targeted analysis did not identify genes that could explain the different flooding tolerance of juvenile and adult plants, we opted for a whole transcriptome approach. For the microarray assay, we based on our previous tolerance tests and chose 24 h dark submergence, as the longest time point before leaf hyperhydricity could be observed in adult individuals. Plants at the same developmental stages maintained for 24 h under continuous darkness were included as controls. As expected, submergence caused profound rearrangements in plant transcriptomes at both ages: out of ∼28000 genes represented by probe sets in the Arabidopsis Gene 1.0 ST Array platform, 1364 were found significantly up- or down-regulated (|Log_2_FC|>2) in common in plants from both ages (Fig. 5a, Table S5a). On the other hand, we found 363 and 232 mRNAs specifically regulated in juvenile or adult plants, respectively (Fig. 5a, Table S5b and c). About half of the genes that were significantly differentially expressed at either age were assigned to the Gene Ontology (GO) categories of response to stress, response to biotic or abiotic stimuli, or other metabolic and cellular processes.

**Fig. 5.**
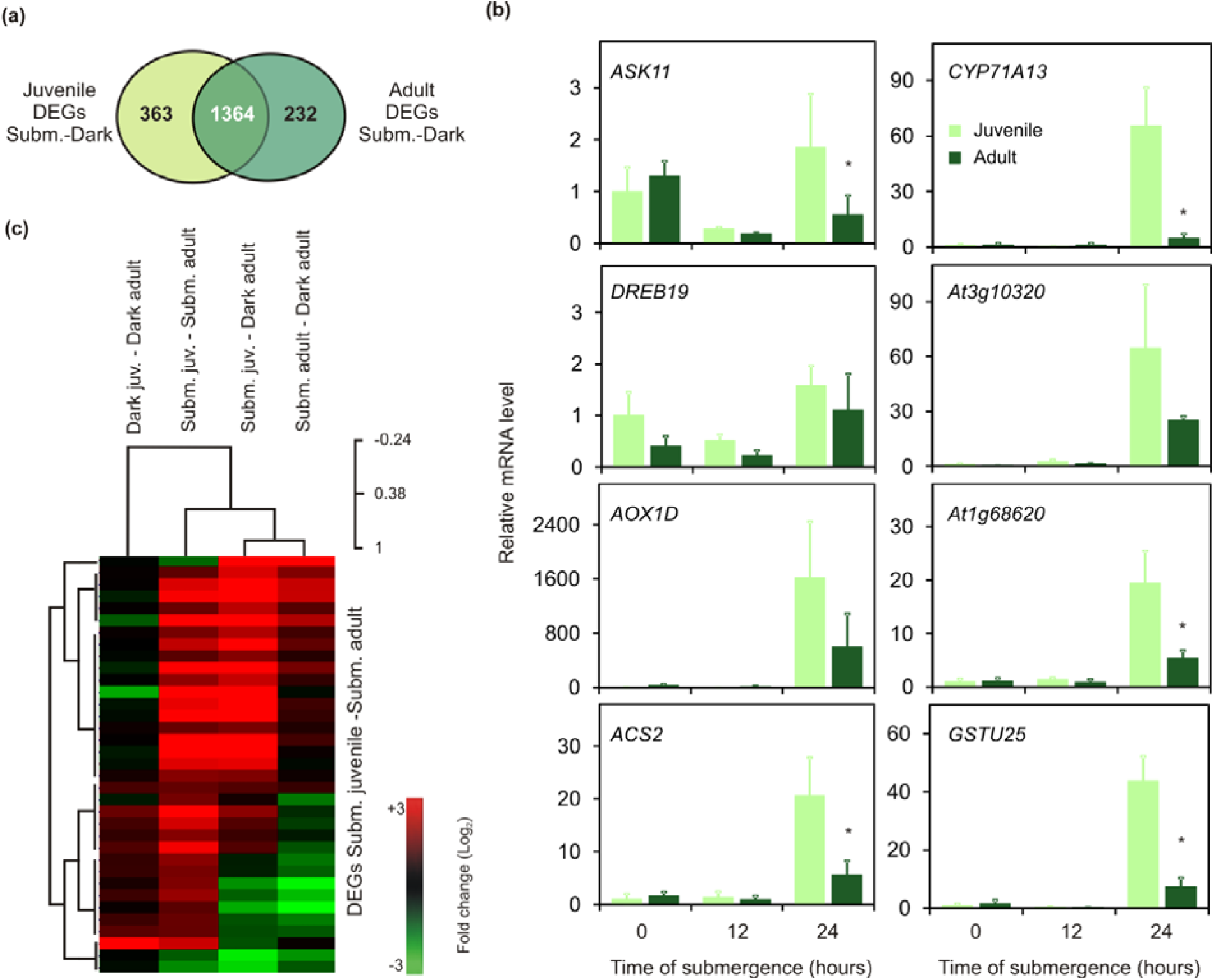
Whole-transcriptome responses of juvenile and adult plants to submergence. **(a)** Venn diagram showing all significant differentially expressed genes (DEGs, |Log_2_FC|>2, adj. p<0.05) found in the microarray comparison between submergence treated (“Subm.”, complete flooding in darkness for 24 h) and control plants (“Dark”, 24 h darkness in air) at each stage of vegetative development. FC, Fold Change. **(b)** Relative expression of a subset of the identified stage-specific DEGs upon submergence, validated by means of a realtime qPCR experiment over a timing of submergence (12 h and 24 h). The expression level recorded in juvenile samples at the beginning of submergence (t=0) was set to 1 for each gene. Data are mean ± S.D. (n=4), asterisks indicate statistically significant difference between adult and juvenile samples at every time point, after t-test (p<0.05). The primer used and the gene AGI codes can be found in Table S1. **(c)** Heatmap of the significant DEGs (|Log_2_FC|>1, adj. p<0.05, stringent selection) between juvenile and adult submergence treated plants, listed in Table 1. Hierarchical clustering of samples across the four possible microarray comparisons (phylogenetic distances are shown on the top right) and of DEGs (left side) was carried out with Multiple Experiment Viewer (Saeed *et al.*, 2003).

We focussed on genes with differential expression in juvenile plants as compared with adult ones under flooding (hereafter, juvenility-specific genes). First, we tested by real time qPCR a selection of eight genes among those with higher expression in submerged juvenile than adult plants (FC ranging from 1.1 to 4.0, 0.03<adj. p<0.1; Table S5a), to confirm their behaviour. Most of them were indeed significantly more up-regulated in juvenile plants as compared to adult ones, after 24 h submergence, confirming the behaviour revealed by the microarray experiment. Instead, we could not observe the same trend after 12 h submergence only (Fig. 5b), suggesting that the observed regulation rather occurred after long-term submergence.

A stringent selection (adj. p<0.05) of the juvenility-specific genes under flooding retrieved 32 genes that showed significantly higher expression (FC>2) in submerged juvenile plants and three that were significantly less expressed (FC<-2) (Table 1 and Table S5d). The 35 genes clearly clustered into two major groups, while clustering together the submergence response data sets within the two ages (Fig. 5c).

To understand the role of these newly identified juvenility-specific genes, we looked at their expression profile and responsiveness in other experimental conditions: a Genevestigator survey over several distinct microarray experiments showed that several of these transcripts respond to abscisic acid (ABA) and antimycin A application, whereas only few of them were affected by low oxygen stresses (Fig. S2 and Table S6). Antimycin A is an inhibitor of the mitochondrial electron transport chain, whose application is known to cause a concomitant drop in ATP levels and the production of reactive oxygen species, similar to what happens under flooding conditions. On the other hand, although drought and ABA-driven response are crucial in the post-submergence phase (Yeung *et al.*, 2018), ABA levels have not been measured in Arabidopsis during submergence, although their ethylene-induced decrease is integral part of tissue elongation during flooding in different species (Hoffmann-Benning & Kende, 1992; Benschop *et al.*, 2005). Considering our data in the light of the existing literature, we therefore hypothesized that either juvenile plants were able to produce higher ABA and/or ROS levels during flooding, or they were more efficient in eliciting the transcriptional response to such stimuli.

### ABA-dependent responses do not contribute to varying submergence tolerance along the vegetative phase change in Arabidopsis

We first focused on ABA synthesis and signalling, and quantified this stress hormone at the beginning and after 12 h and 24 h of submergence using an ELISA immunodetection assay. The only observable difference was a higher ABA content in adult plants prior to treatment (Fig. 6a). In agreement with previous reports, we then observed a decrease in the level of this hormone upon submergence, which occurred to the same extent at both developmental stages (Fig. 6a). These pieces of evidence led us to discard the hypothesis that higher ABA content in juvenile plants could explain the difference in transcriptional response to flooding.

**Fig. 6.**
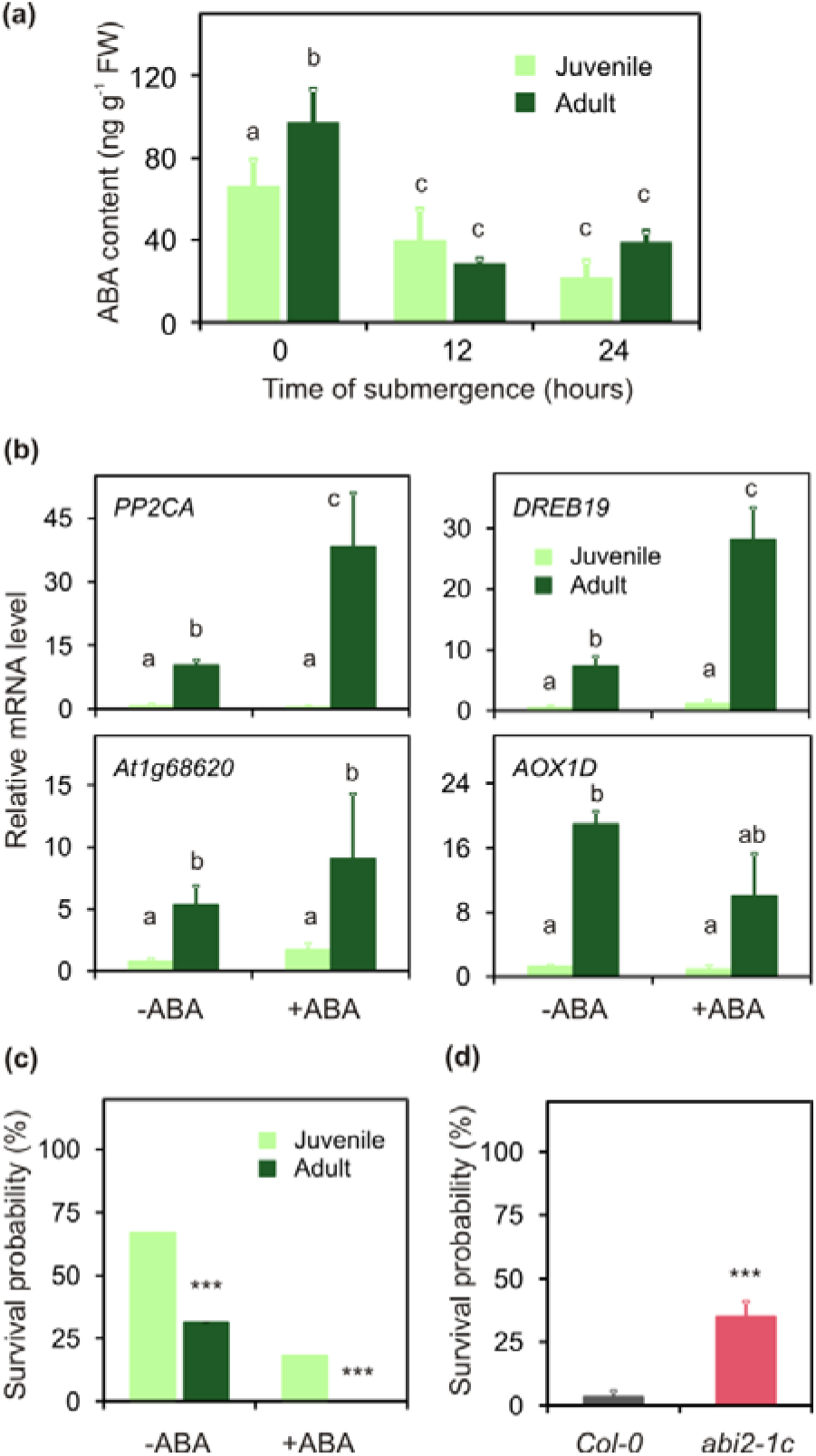
Impact of ABA levels and signalling on flooding tolerance across developmental stages. **(a)** ABA content in the rosette of two and three week-old plants at different time points of submergence or prior to treatment. Letters mark statistically significant differences after two-way ANOVA analysis and Holm-Sidak post hoc test (p<0.05). **(b)** Relative mRNA levels of ABA-responsive genes in juvenile and adult plants treated with 10 μM ABA or water, for 3 h. The expression level recorded in juvenile samples at the beginning of the treatment was set to 1 for each gene. Data are mean ± S.D. (n=4), different letters indicate statistically significant differences between adult and juvenile samples after two-way ANOVA (p<0.05) and Tukey’s post-hoc test. **(c)** Flooding tolerance of juvenile and adult plants treated with 0 or 10 μM ABA 3 hours before a 48 hour-long submergence. **(d)** Tolerance of wild type and *abi2-1c* adult plants (n=15) subjected to 72 h dark submergence. Data are presented as mean ± S.E (n=4). Asterisks indicate statistically significant differences (p<0.01) as assessed by Kaplan-Meier analysis.

To evaluate whether the juvenile tolerance could arise from some enhanced sensitivity to the hormone, we thus moved on to compare plant responsiveness to ABA. Using the dedicated Genevestigator tool, we selected five ABA-responsive markers from publicly available experiments: *PP2CA (At3g11410), DREB19 (At2g38340)*, the alpha/beta hydrolase *At1g68620, AOX1D (At1g32350)*, and *ACS2 (At1g01480)*. Their expression was analysed by means of qPCR in adult and juvenile plants treated with 10 μM ABA for 3 h. Apart from *ACS2*, which was not induced by the treatment (not shown), the remaining four genes displayed overall higher basal expression and stronger induction by ABA (with the exception of *AOX1D*) in adult plants, suggesting that juvenile plants are not likely to be more sensitive to the hormone under unstressed conditions (Fig 6b). Not only the more submergence-tolerant juvenile plants showed no differential responses to ABA, but also ABA signalling proved unable to exert a positive impact on flooding tolerance in plants of different ages. On one hand, ABA pre-treatment 3 hours before submergence decreased survival probability at both developmental stages (Fig. 6c).

The results obtained in the analyses described above led us to dismiss the hypothesis of a prominent role by ABA signalling in enhancing tolerance in juvenile plants. On the other hand, these results could suggest that ABA signalling contributes to plant sensitivity to submergence. We tested this by means of *abi2-1* mutant plants, which are ABA insensitive. Indeed, adult *abi2-1* plants displayed significantly better tolerance to submergence than the wild type (Fig. 6d). At this point, determined to pursue the initial question about the differential expression of the identified set of juvenile-specific genes, we moved on to evaluate the contribution of ROS synthesis, perception and downstream signalling.

### The age-dependent sensitivity to submergence is due to a differential activity of the ROS-regulated transcription factor ANAC017

To substantiate the connection between higher expression of ROS-related genes in juvenile plants (Fig. S2b) and better flooding tolerance of plants at this stage, we first tested whether adult plants can be primed to better endure submergence by application of a low dose of antimycin A (100 nM) prior to stress. In this sense, antimycin A pre-treatment proved indeed effective for adult plants (Fig. 7a and b), while it did not produce any effect on two week old plants (Fig. 7b). This suggests that promotion of juvenile-specific gene expression in older plants can enable them to better cope with flooding conditions.

**Fig. 7.**
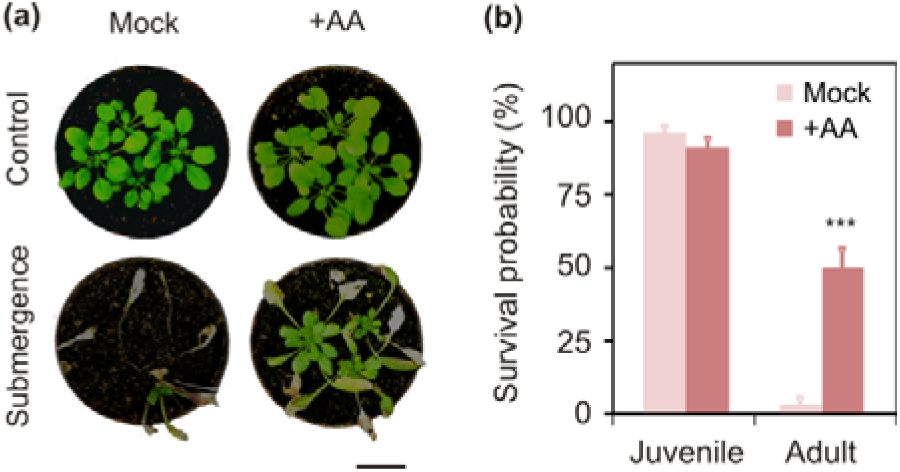
Antimycin A treatment improved submergence tolerance in adult but not juvenile plants. **(a)** Representative phenotype of adult plants portrayed one week after recovery, following dark submergence stress. Plants were sprayed with 100 nM antimycin A (+AA) or 0.001% ethanol solutions (“mock”) 6 hours prior to 24 hour-long flooding (“submergence”). Scale bar, 2 cm. **(b)** Flooding tolerance of AA-treated plants of both juvenile and adult ages (n=20). Data are presented as mean ± S.E. (n=4). Asterisks indicate statistically significance at p<0.05 with Kaplan-Meier analysis.

Higher expression levels of ROS-related genes in juvenile plants might be a consequence of better plant ability to activate ROS signalling under submergence, and thereby downstream tolerance strategies, at that stage. However, a qualitative assessment of H_2_O_2_ accumulation by diaminobenzidine (DAB) staining after 24 h submergence did not show remarkable differences between juvenile and adult plants (Fig. S3a); at both ages, ROS production was stimulated by dark submergence and, to a lower extent, by extended prolonged darkness conditions (Fig. S3b). Therefore, we favoured the alternative explanation, by which juvenile plants might be especially sensitive to ROS-related signals produced under flooding.

The inhibition of the mETC and consequent ROS production in Arabidopsis has been shown to activate a number of genes involved in the attenuation of oxidative stress by means of a subgroup of NAC transcription factors (Ng et al., 2013). These proteins, which include ANAC013 (At1g32870), 16 (At1g34180), and 17 (At1g34190), are characterized by a transmembrane region at the C-terminus that maintains them in the endoplasmic reticulum (ER) (De Clercq et al., 2013a; Ng et al., 2013). Upon oxidative stress, a not-yet identified signal of mitochondrial origin has promotes the proteolytic cleavage of the transmembrane domain of ANAC013 and 17 via rhomboid proteases, thereby allowing the cleaved soluble polypeptide to re-localize to the nucleus. We therefore investigated the involvement of these transcription factors in the superior tolerance of juvenile plants to flooding. Within the broad NAC family, by blast analysis we could identify five members with high sequence similarity to ANAC017 and a putative transmembrane region at the C-terminus. ANAC053 (At3g10500) and 78 (At5g04410), which formed a separate clade from ANAC013, 16 and 17 (Fig. 8a), have been recently reported to regulate the response to proteotoxic stress (Gladman *et al.*, 2016). Limited to the members of the latter clade, interrogation of public gene expression datasets did not show significant differences at the transcriptional level between leaf samples from young or developed plants at the pre-flowering stage (Fig. S4). Based on these observations, we decided to focus on ANAC017 as the most highly expressed member of the subgroup in rosette tissues (Fig. S4).

**Fig. 8.**
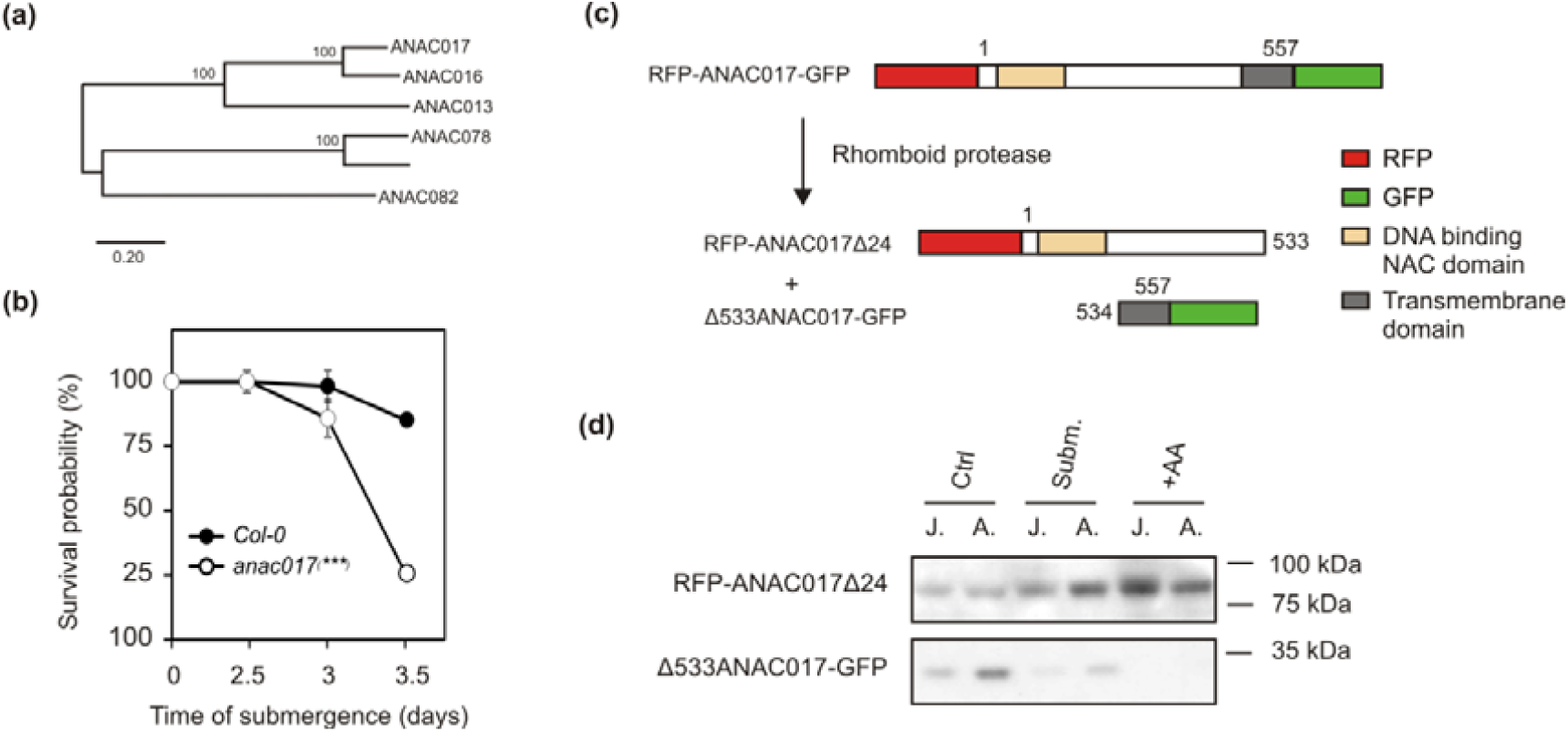
Involvement of ANAC017 in stage-specific responses to flooding. **(a)** Phylogenetic relationships among the NAC transcription factor family proteins with highest sequence similarity to the ANAC017 member. Numbers indicate phylogenetic distances calculated with Mega10 with 500 bootstrap value. ANAC082 (At5g09330), a family member lacking any transmembrane domain, was employed in the analysis to root the phylogenetic tree. **(b)** Submergence tolerance in two week old *anac017* mutant plants (n=15), as compared with wild type. Data are presented as mean ± S.E. (n=4). Asterisks indicate significant difference (p<0.001) with Kaplan-Meier analysis. **(c)** Schematic representation of a reporter construct for the ANAC017 protein, introduced in plants, and its hypothetical post-transcriptional regulation. In the presence of an oxidative stress, the protein is expected to be cleaved by rhomboid protease family enzymes. In the transgenic plants, the fate of the N-terminal and C-terminal halves can be followed thanks to the RFP and GFP tags fused to the full length ANAC017 sequence (557 amino acid-long), as depicted. **(d)** Immunoblotting of juvenile (“J.”) or adult plant (“A.”) total protein extracts from rosette leaves with either an RFP-specific antibody (above) or a GFP-specific one (below). Plants were kept under submergence (“Subm.”) or extended darkness (“Ctrl”) for 24 h, or sprayed with 10 µM antimycin A 6 h before harvest. The observed bands match the expected size of an RFP-ANAC017Δ24 peptide (calculated MW≈70 kDa) and a Δ533ANAC017-GFP peptide (calculated MW≈30 kDa). Full blots are displayed in Fig. S7.

When challenged with submergence, a homozygous knock-out ANAC017 mutant (*anac017-1)*, exhibited significantly lower survival at the juvenile stage, as compared to wild type plants of the same age (Fig. 8b), providing evidence for the involvement of this transcription factor in flooding tolerance. Although a contribution by ANAC017 homologs to plant performance under submergence cannot be ruled out, the extent of reduction in flooding tolerance we observed in *anac017* supports the hypothesis of a major role of this transcription tolerance in the response of juvenile Arabidopsis plants during the stress. Remarkably, most of the juvenile-specific genes that are responsive to antimycin A appeared to be less up-regulated (or sometimes even down-regulated) in juvenile *anac017* mutant plants (Fig. S2b). A subset of these genes also proved to be less induced upon submergence in juvenile *anac017* plants (Fig. S5).

Next, we tested whether ANAC017 is differentially regulated at the post-transcriptional level in juvenile and adult plants. To this purpose, we cloned its full length coding sequence and fused it to an N-terminal RFP sequence and a C-terminal GFP, which should enable us to follow the fate of the two protein halves subsequently to cleavage (Fig. 8c). This construct was transformed in Arabidopsis Col-0 plants under control of the Arabidopsis *UBQ10* promoter. Similar to what has been reported before for different ANAC017 over-expressors (Meng *et al.*, 2019), adult *pUBQ10:RFP-ANAC017-GFP* plants had altered phenotype with shorter petioles and slightly adaxialised leaves at the adult stage (Fig. S6), although they were indistinguishable from the wild type at the juvenile stage. We investigated ANAC017 cleavage upon mitochondrial stress across the two different developmental stages, in plants experiencing submergence (24 h), extended darkness (24 h), or antimycin A treatment (10 µM, 6 h). Immunodetection of RFP and GFP in total rosette protein extracts showed fragments of about 70 kDa and 30 kDa, respectively, corresponding to the calculated size of the hypothetical cleavage products, RFP-ANAC017Δ24 (retaining an ANAC017_1-533_ fragment) and Δ533ANAC017-GFP (ANAC017_534-557_) (Fig. 8d). In addition to these, the anti-RFP antibody could detect a 35 KDa band, most likely corresponding to an N-terminal fragment of ANAC017, and one around 100 KDa, which we attributed to the uncleaved version of the transcription factor (Fig. 8d). The nuclear targeted fragment RFP-ANAC017Δ24 accumulated at higher levels both under flooding and antimycin A treatments than in control conditions, while the ER-localized fragment strongly decreased upon stress treatment independently of the age considered (Fig. 8d). Juvenile plants accumulated more RFP-ANAC017Δ24 when treated with antimycin A, but not under submergence. Considered together, these results suggested that different ANAC017 proteolysis upon submergence does not account for the ROS-mediated superior tolerance of juvenile plants.

### Stage-specific chromatin modifications occur in vegetative Arabidopsis tissues at juvenile-specific loci

Having excluded the occurrence of transcriptional or post-transcriptional regulation on ANAC017 in the developmental stages under investigation, we pointed at the epigenetic status of the target DNA loci and investigated some chromatin modifications in the same selected juvenile-specific genes described above. To this end, we kept the analysis focussed on the same set of genes evaluated in Fig. 5b, as interrogation of public datasets confirmed their ANAC017-dependent upregulation in response to mitochondrial stress (Fig. 9a). A possible mechanism underlying the differences observed in terms of mRNA accumulation (Fig. 5b) might consist in distinct age-specific methylation patterns of the genomic DNA, able to affect RNA polymerase activity. We exploited the methylation-sensitive restriction enzyme-qPCR technique (MSRE-qPCR) (Hashimoto *et al.*, 2007) to evaluate cytosine methylation of particular HpaII and SalI sites found on the target loci (Table S2). With the exception of *CYP71A13* and *GSTU25*, the analysis revealed a trend for higher methylation of the target loci in juvenile samples (Fig. 9b).

**Fig. 9.**
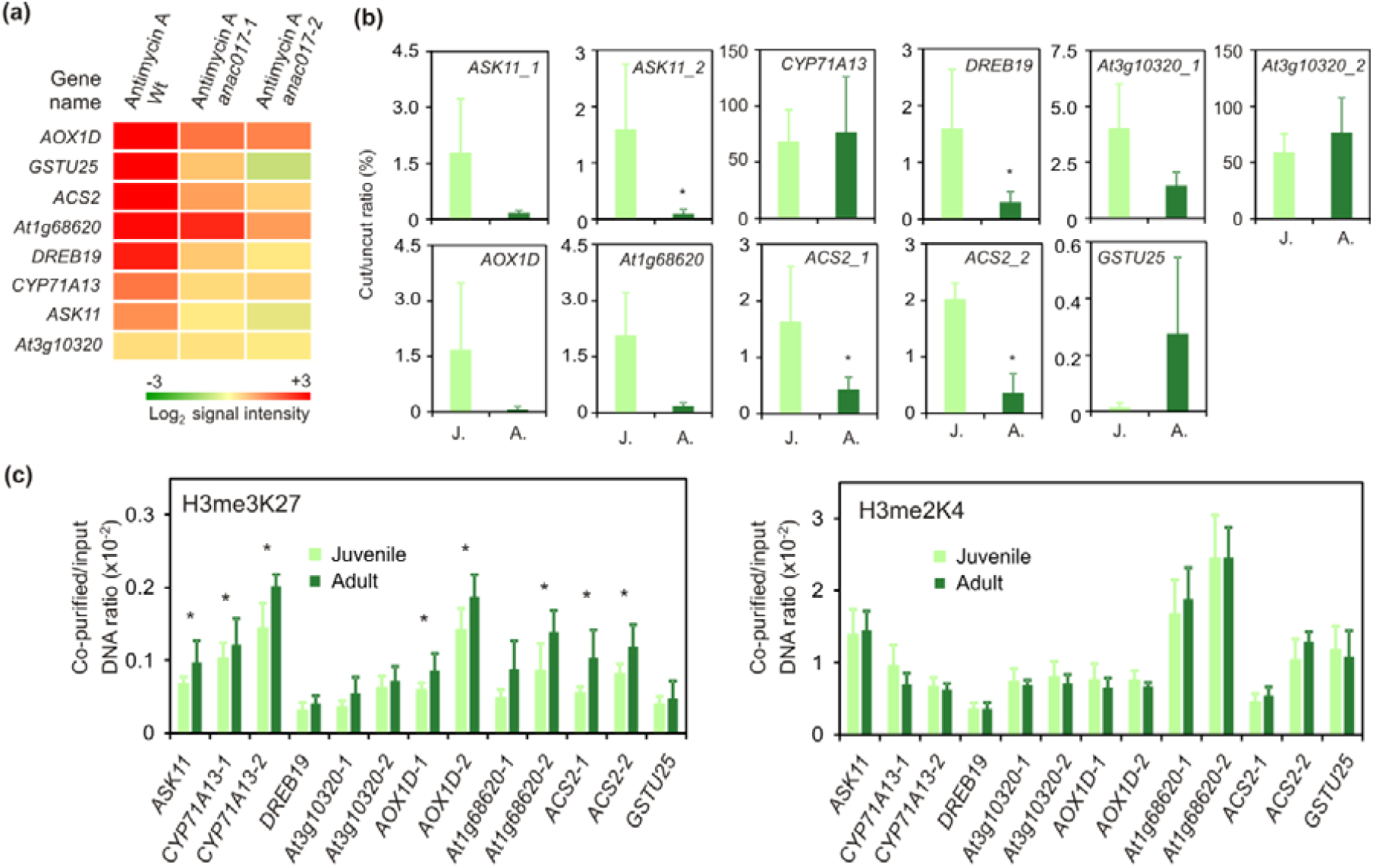
Evaluation of promoter DNA methylation and chromatin accessibility at selected juvenile-specific loci. **(a)** Expression patterns of the juvenile-specific genes also shown in Figure 5b, extracted with Genevestigator from public microarray datasets, showing gene response to antimycin A in two independent *anac017* T-DNA mutant lines and in a background genotype (“Wt”). The dataset used are specified in Table S6. **(b)** MSRE-qPCR results on the target genes. Higher ratio is suggestive of higher methylation at the target DNA sequence. Genomic DNA extracted from juvenile (J.) or adult (A.) leaf samples was digested using the methylation sensitive enzymes HspII or SalI, according to the information provided in Table S2, or mock-digested. Cut or uncut DNA was amplified using the primers listed in Table S2. Data are mean ± S.D. (n=4). Asterisks indicate statistically significant differences after t-test (p<0.05). **(b)** H3K27 tri-methylation and HK4 di-methylation, revealed my ChIP-qPCR. Related information can be found in Table S3. Data are mean ± S.D. (n=5). Asterisks indicate statistically significant differences after t-test (p<0.05).

In another scenario, juvenile and adult plants might differ in terms of chromatin accessibility to ANAC017, after its activation by mitochondrial stress signalling, and to other regulatory factors. We thus analysed a histone 3 modification of known to play a repressive role in the octamer, K27 trimethylation (H3me3K27), and found that five of the gene promoters evaluated, namely *ASK11, CYP71A13, AOX1D, At1g68620* and *ACS2*, showed significantly higher enrichment in adult plants in comparison with juvenile ones, in at least one analyzed region (Fig. 9c). This observation hints at a lower extent of chromatin repression at the target loci in juvenile plants, which might facilitate the induction of such genes upon the onset of submergence. A second epigenetic marker of repression, H3K4 dimethylation (H3me2K4, (Liu *et al.*, 2019)), showed instead no significant differences in our conditions (Fig. 9c).

## Discussion

Plants, including Arabidopsis, shift through developmental phases integrating cues from the environment and by modulating genetic programmes. To the best of our knowledge, this study represents the first attempt to investigate submergence tolerance in Arabidopsis at different stages of plant growth. We consistently observed higher tolerance of juvenile plants in comparison to older ones (Fig. 1 and 2). The transition from juvenility to adulthood is accompanied by an increase in the level of SPL proteins. Inversely, the elevation of SPL protein correlates with the down regulation of the miR156 transcript, which acts as a negative regulator of *SPL* genes (Wu et al., 2009; Huijser and Schmid, 2011). Here, we confirmed that adult plants grown in our experimental conditions increase significantly the expression of *SPL3* and *SPL4* as compared to juvenile plants (Fig. 3b). Surprisingly, instead, artificial procrastination of adulthood by miR156 overexpression did not improve the tolerance to submergence (Fig. 3c and d), indicating that either parallel signalling pathways, associated with juvenility to adulthood transition, are responsible for this phenomenon, or that ectopic expression of the miRNA impaired the plant’s ability to respond to flooding.

The molecular response to submergence broadly overlapped between juvenile and adult plants, and corresponded to the one that we previously reported for four week-old plants (Giuntoli et al. 2017). The typical hypoxia-marker genes were strongly induced after 12 and 24 h of submergence, as well as transcripts coding for proteins related to ROS homeostasis, protein stabilization and defense mechanisms (Fig. 4). Juvenile plants exhibited faster and stronger induction of *ADH*, suggestive of enhanced fermentation, while adult plants showed the highest induction of starvation-related genes *DIN6* and *TPS8.* This observation was expected, since carbon availability has been shown to affect survival to submergence in Arabidopsis by modulating the magnitude with which ERF-VII transcription factors regulate the expression of fermentative genes (Loreti et al. 2018). Indeed, juvenile plants have smaller leaves than adult ones, with higher carbon reserves, including sucrose and soluble sugars (Durand *et al.*, 2018). Thus, we speculate that the superior tolerance to submergence in juvenile plants is associated to the promptness and extent of the metabolic and molecular responses activated at this stage.

Under submergence conditions, the amount of transcripts identified as differentially expressed between juvenile and adult individuals was relatively small (Table 1). Several of them have been reported to respond to treatments that produce reactive oxygen species or activate the ABA signalling pathway. A real time qPCR survey, carried out on a subset of these genes, revealed that they are upregulated after 24 h but not 12 h of submergence, indicating that this response is activated late during submergence (Fig. 5b).

ABA has been investigated as a hormone potentially involved in submergence responses in several plant species, including both dicots and monocots (Hook *et al.*, 1988; Hoson *et al.*, 1993; Hurng *et al.*, 1994; Lee *et al.*, 2009). Consistent with previous reports, our data confirmed that ABA levels decreased upon submergence at both developmental stages (Fig. 6a). Adult plants showed stronger ABA-responsiveness under non-flooded conditions (Fig. 6b), suggesting that other signals beyond ABA are involved in the upregulation of the subset of juvenile-specific genes under submergence. ABA is actually more likely to play a negative role in the adaptation of plants to submergence since ABA pre-treatment reduced submergence survival at both ages (Fig. 6c). Accordingly, an ABA-insensitive mutant (*abi2-1*) exhibited increased flooding tolerance in adult plants (Fig. 6d). Taken together, our data excluded increased ABA levels or responsiveness as determinants of superior tolerance to submergence in juvenile Arabidopsis plants. Instead, they suggested that plant sensitivity to submergence entails functional ABA signalling.

Instead, we could link this feature to the impairment of mitochondrial activity. Treatment with antimycin A, which inhibits mtETC complex III and consequently leads to the production of ROS, was reported to cause upregulation of one third of the juvenile-specific genes (Fig. S2b). These were mainly genes related to ROS homeostasis, mitochondrial functioning and cell wall remodelling, which potentially improve survival under stress conditions. Indeed, pre-treatment of adult plants with antimycin A significantly improved flooding survival in adult plants but not juvenile ones. We speculated that either juvenile plants at this developmental stage are especially sensitive to ROS-related signals produced under submergence conditions or that they actually are more promptly able to produce this signal when the stress occurs. We could not detect differences between juvenile and adult plants after DAB staining for hydrogen peroxide under control or stress conditions (Fig. S3).

A group of ER-bound NAC transcription factors has been reported as main regulators to the nuclear response to mitochondrial dysfunction (De Clercq *et al.*, 2013; Ng *et al.*, 2013; Meng *et al.*, 2019). Mitochondrial ROS production has been proposed to lead to the release of N-terminal fragments of these factors, by action of rhomboid proteases, into the nucleus, where they act as transcriptional regulators. Among them, ANAC017 was abundantly expressed in both juvenile and adult plants (Fig. S4) and T-DNA mediated inactivation of *ANAC017* reduced significantly survival under submergence condition compared with the wild type (Fig. 8b). By exploiting dual fluorescent tagging of the protein (Fig. 8c), we observed constitutive endoproteolytic cleavage of ANAC017 under control conditions at both developmental stages considered (Fig. 8d). We also observed accumulation of the N-terminal fragment in case of submergence and antimycin A treatment while the C-terminal fragment decreased after both treatments (Fig. 8d). This observation would invoke additional proteostatic mechanisms acting upon this transcription factor after its proteolytic cleavage. In addition, ANAC017 transgenic plants developed their phenotype upon aging. We could observe no difference in phenotype at the juvenile stage but the two genotypes became distinguishable, as reported by (Meng *et al.*, 2019), after the third week (Fig. S6). This could correlate with an increased abundance of active, nuclear localized, ANAC017 when plant development proceeds with age. The transcriptional activity of ANAC013 and ANAC017 has been recently shown to be regulated by the nuclear protein RADICAL-INDUCED CELL DEATH1 (RCD1) (Shapiguzov *et al.*, 2019) and the relevance of such interaction might be investigated in the future in the context of flooding stresses.

While the abundance of the N-terminal ANAC017 fragment was equal between juvenile and adult plants (Fig. 8d), we observed a difference in the accessibility of its target genes at the chromatin level. Indeed, we could measure enhanced trimethylation of H3K27, a marker of gene repression (Mondal *et al.*, 2016; Pan *et al.*, 2018), in adult plants (Fig. 9c), when considering promoters of representative ANAC017 target genes, identified among the juvenile-specific genes (Fig. 9a) upon survey of publicly available transcriptomic data (Fig. 9a). The same genomic regions modifications were characterized by lower DNA methylation, in line with what has been described for animal models (Manzo *et al.*, 2017). Promotion of a heterochromatic context in specific stress-related genes in adult plants might be interpreted as an adaptive strategy to limit responsiveness, and its high energy costs, in plants destined to reproductive development.

To conclude, in the same genetic background, juvenile Arabidopsis plants showed enhanced survival than adult plants under submergence conditions. We showed that this tolerance mechanism is independent of the core-anaerobic response (Fig. 4) and rather relies on NAC transcription factors that mediate retrograde stress signalling (Fig. 8b and 9). The differential response between developmental stages seems to originate from the chromatin status of the loci targeted by ANAC017, rather than from its direct regulation by the stress. If confirmed in crop species, this finding might help breeding programs and farming practice to tailor strategies on the specific developmental stage at which submergence-related stresses are experienced.

## Acknowledgements

This study was financially supported by the Scuola Superiore Sant’Anna and the PRIN project (Oxygen dependent control of plant development), funded by the Italian Ministry of University and Research. LTB was financially supported by the PhD programme in Agrobiodiversity.

## Author contributions

FL, LTB and BG designed the experiments. LTB carried out survival analyses, tissue stainings, gene-targetted expression analyses, gene cloning and the production of transgenic plants, in addition to the statistical analyses associated with the above-mentioned experiments. BG carried out immunoblot and MSRE-qPCR analyses. AT quantified ABA levels. FMG analysed the microarray data. VS carried out Chip-PCR analyses. FL and PP secured funding to support the study. FL and BG wrote the manuscript, with inputs by PP. All authors read and approved the manuscript.

## References

Bailey-Serres J, Colmer TD. 2014. Plant tolerance of flooding stress – recent advances. Plant, Cell & Environment 37: 2211–2215.

Bechtold U, Field B. 2018. Molecular mechanisms controlling plant growth during abiotic stress. Journal of Experimental Botany 69: 2753–2758.

Benschop JJ, Jackson MB, Gühl K, Vreeburg RAM, Croker SJ, Peeters AJM, Voesenek LACJ. 2005. Contrasting interactions between ethylene and abscisic acid in Rumex species differing in submergence tolerance. Plant Journal 44: 756–768.

Boyes DC. 2001. Growth stage-based phenotypic analysis of arabidopsis: a model for high throughput functional genomics in plants. The Plant Cell 13: 1499–1510.

De Clercq I, Vermeirssen V, Van Aken O, Vandepoele K, Murcha MW, Law SR, Inzé A, Ng S, Ivanova A, Rombaut D, et al. 2013. The membrane-bound NAC transcription factor ANAC013 functions in mitochondrial retrograde regulation of the oxidative stress response in Arabidopsis. The Plant Cell 25: 3472–3490.

Daudi A, O’Brien JA. 2012. Detection of hydrogen peroxide by DAB staining in Arabidopsis leaves. Bio-protocol 2: e263.

van Dongen JT, Licausi F. 2015. Oxygen sensing and signaling. Annual Review of Plant Biology 66: 345–346.

Durand M, Mainson D, Porcheron B, Maurousset L, Lemoine R, Pourtau N. 2018. Carbon source–sink relationship in Arabidopsis thaliana: the role of sucrose transporters. Planta 247: 587–611.

Gibbs DJ, Lee SC, Md Isa N, Gramuglia S, Fukao T, Bassel GW, Correia CS, Corbineau F, Theodoulou FL, Bailey-Serres J, et al. 2011. Homeostatic response to hypoxia is regulated by the N-end rule pathway in plants. Nature 479: 415–418.

Giorgi FM, Bolger AM, Lohse M, Usadel B. 2010. Algorithm-driven Artifacts in median polish summarization of Microarray data. BMC Bioinformatics 11: 553.

Giuntoli B, Perata P. 2017. Group VII Ethylene Response Factors in Arabidopsis: regulation and physiological roles. Plant Physiology 176: 1143–1155.

Gladman NP, Marshall RS, Lee KH, Vierstra RD. 2016. The proteasome stress regulon is controlled by a pair of NAC transcription factors in arabidopsis. Plant Cell 28: 1279–1296.

Gonzali S, Loreti E, Cardarelli F, Novi G, Parlanti S, Pucciariello C, Bassolino L, Banti V, Licausi F, Perata P. 2015. Universal stress protein HRU1 mediates ROS homeostasis under anoxia. Nature Plants 1: 15151.

Grefen C, Donald N, Hashimoto K, Kudla J, Schumacher K, Blatt MR. 2010. A ubiquitin-10 promoter-based vector set for fluorescent protein tagging facilitates temporal stability and native protein distribution in transient and stable expression studies. Plant Journal 64: 355–365.

Hashimoto K, Kokubun S, Itoi E, Roach HI. 2007. Improved quantification of DNA methylation using methylation-sensitive restriction enzymes and real-time PCR. Epigenetics 2: 86–91.

Hildebrand F. 1875. Ueber die Jungendzustände solcher Pflanzen, welche im Alter vom vegetativen Charakter ihrer Verwandten abweichen. Flora 21: 321–330.

Hoffmann-Benning S, Kende H. 1992. On the role of abscisic acid and gibberellin in the regulation of growth in rice. Plant Physiology 99: 1156–1161.

Hook DD, McKee WH, Smith HK, Gregory J, Burrell VG, DeVoe MR, Sojka RE, Gilbert S, Banks R, Stolzy LH, et al. 1988. Involvement of the hormones ethylene and abscisic acid in some adaptive responses of plants to submergence, soil waterlogging and oxygen shortage. The ecology and management of wetlands, Springer, NY, p. 373–382.

Hoson T, Masuda Y, Pilet PE. 1993. Abscisic acid content in air- and water-grown rice coleoptiles. Journal of Plant Physiology 142: 593–596.

Hruz T, Laule O, Szabo G, Wessendorp F, Bleuler S, Oertle L, Widmayer P, Gruissem W, Zimmermann P. 2008. Genevestigator V3: A Reference Expression Database for the Meta-Analysis of Transcriptomes. Advances in Bioinformatics 2008: 1–5.

Huijser P, Schmid M. 2011. The control of developmental phase transitions in plants. Development 138: 4117–4129.

Hurng WP, Lur HS, Liao CK, Kao CH. 1994. Role of abscisic acid, ethylene and polyamines in flooding-promoted senescence of tobacco leaves. Journal of Plant Physiology 143: 102–105.

James SA, Bell DT. 2001. Leaf morphological and anatomical characteristics of heteroblastic Eucalyptus globulus ssp. Globulus (Myrtaceae). Australian Journal of Botany 49: 259–269.

Kanojia A, Dijkwel PP. 2018. Abiotic stress responses are governed by reactive oxygen species and age. Annual Plant Reviews online: 1–32.

Karimi M, Inzé D, Depicker A. 2002. GATEWAY^TM^ vectors for Agrobacterium-mediated plant transformation. Trends in Plant Science 7: 193–195.

Lee B, Yu S, Jackson D. 2009. Control of plant architecture: the role of phyllotaxy and plastochron. Journal of Plant Biology 52: 277–282.

Licausi F, Kosmacz M, Weits DA, Giuntoli B, Giorgi FM, Voesenek LACJ, Perata P, Van Dongen JT. 2011. Oxygen sensing in plants is mediated by an N-end rule pathway for protein destabilization. Nature 479: 419–422.

Lim CCK, Krebs SL, Arora R. 2014. Cold hardiness increases with age in juvenile rhododendron populations. Frontiers in Plant Science 5: 542.

Liu Y, Liu K, Yin L, Yu Y, Qi J, Shen WH, Zhu J, Zhang Y, Dong A. 2019. H3K4me2 functions as a repressive epigenetic mark in plants. Epigenetics and Chromatin 12: 40.

Livak KJ, Schmittgen TD. 2001. Analysis of relative gene expression data using real-time quantitative PCR and the 2(-Delta Delta C(T)) Method. Methods 25: 402–8.

Loreti E, van Veen H, Perata P. 2016. Plant responses to flooding stress. Current Opinion in Plant Biology 33: 64–71.

Manzo M, Wirz J, Ambrosi C, Villaseñor R, Roschitzki B, Baubec T. 2017. Isoform-specific localization of DNMT3A regulates DNA methylation fidelity at bivalent CpG islands. The EMBO Journal 36: 3421–3434.

Marias DE, Meinzer FC, Still C. 2017. Impacts of leaf age and heat stress duration on photosynthetic gas exchange and foliar nonstructural carbohydrates in Coffea arabica. Ecology and Evolution 7: 1297–1310.

Matsoukas IG, Massiah AJ, Thomas B. 2013. Starch metabolism and antiflorigenic signals modulate the juvenile-to-adult phase transition in Arabidopsis. Plant, Cell and Environment 36: 1802–1811.

Meng X, Li L, Clercq I De, Narsai R, Xu Y, Hartmann A, Claros DL, Custovic E, Lewsey MG, Whelan J, et al. 2019. ANAC017 coordinates organellar functions and stress responses by reprogramming retrograde signaling. Plant Physiology 180: 634–653.

Mithran M, Paparelli E, Novi G, Perata P, Loreti E. 2014. Analysis of the role of the pyruvate decarboxylase gene family in Arabidopsis thaliana under low-oxygen conditions. Plant Biology 16: 28–34.

Mondal S, Go YS, Lee SS, Chung BY, Kim JH. 2016. Characterization of histone modifications associated with DNA damage repair genes upon exposure to gamma rays in Arabidopsis seedlings. Journal of Radiation Research 57: 646–654.

Ng S, Ivanova A, Duncan O, Law SR, Van Aken O, De Clercq I, Wang Y, Carrie C, Xu L, Kmiec B, et al. 2013. A membrane-bound NAC transcription factor, ANAC017, mediates mitochondrial retrograde signaling in Arabidopsis. Plant Cell 25: 3450–3471.

Pan MR, Hsu MC, Chen LT, Hung WC. 2018. Orchestration of H3K27 methylation: mechanisms and therapeutic implication. Cellular and Molecular Life Sciences 75: 209–223.

Poethig RS. 2013. Vegetative phase change and shoot maturation in plants. Current Topics in Developmental Biology 105: 125–152.

Ritchie ME, Phipson B, Wu D, Hu Y, Law CW, Shi W, Smyth GK. 2015. Limma powers differential expression analyses for RNA-sequencing and microarray studies. Nucleic Acids Research 43: e47.

Rueden CT, Schindelin J, Hiner MC, DeZonia BE, Walter AE, Arena ET, Eliceiri KW. 2017. ImageJ2: ImageJ for the next generation of scientific image data. BMC Bioinformatics 18: 529.

Saeed AI, Sharov V, White J, Li J, Liang W, Bhagabati N, Braisted J, Klapa M, Currier T, Thiagarajan M, et al. 2003. TM4: A free, open-source system for microarray data management and analysis. BioTechniques 34: 374–378.

Shapiguzov A, Vainonen JP, Hunter K, Tossavainen H, Tiwari A, Järvi S, Hellman M, Aarabi F, Alseekh S, Wybouw B, et al. 2019. Arabidopsis RCD1 coordinates chloroplast and mitochondrial functions through interaction with ANAC transcription factors. eLife 8: pii: e43284.

Telfer A, Bollman KM, Poethig RS. 1997. Phase change and the regulation of trichome distribution in *Arabidopsis thaliana*. Development 124: 645–654.

Tsukaya H, Shoda K, Kim GT, Uchimiya H. 2000. Heteroblasty in *Arabidopsis thaliana* (L.) Heynh. Planta 210: 536–542.

Voesenek LACJ, Bailey-Serres J. 2015. Flood adaptive traits and processes: an overview. New Phytologist 206: 57–73.

Wang JW, Czech B, Weigel D. 2009. miR156-Regulated SPL transcription factors define an endogenous flowering pathway in *Arabidopsis thaliana*. Cell 138: 738–749.

Weits DA, Giuntoli B, Kosmacz M, Parlanti S, Hubberten HM, Riegler H, Hoefgen R, Perata P, Van Dongen JT, Licausi F. 2014. Plant cysteine oxidases control the oxygen-dependent branch of the N-end-rule pathway. Nature Communications 5: 3425.

White MD, Klecker M, Hopkinson RJ, Weits DA, Mueller C, Naumann C, O’Neill R, Wickens J, Yang J, Brooks-Bartlett JC, et al. 2017. Plant cysteine oxidases are dioxygenases that directly enable arginyl transferase-catalysed arginylation of N-end rule targets. Nature Communications 8: 14690.

Wingler A. 2018. Transitioning to the next phase: The role of sugar signaling throughout the plant life cycle. Plant Physiology 176: 1075–1084.

Woo D-H, Park H-Y, Kang IS, Lee S-Y, Moon BY, Lee CB, Moon Y-H. 2011. Arabidopsis lenc1 mutant displays reduced ABA accumulation by low AtNCED3 expression under osmotic stress. Journal of plant physiology 168: 140–147.

Yeung E, Bailey-Serres J, Sasidharan R. 2019. After the deluge: plant revival post-flooding. Trends in Plant Science 24: 443–454.

Yeung E, van Veen H, Vashisht D, Paiva ALS, Hummel M, Rankenberg T, Steffens B, Steffen-Heins A, Sauter M, de Vries M, et al. 2018. A stress recovery signaling network for enhanced flooding tolerance in *Arabidopsis thaliana*. Proceedings of the National Academy of Sciences of the United States of America 115: E6085–E6094.

Yu S, Li C, Zhou CM, Zhang TQ, Lian H, Sun Y, Wu J, Huang J, Wang G, Wang JW. 2013. Sugar is an endogenous cue for juvenile-to-adult phase transition in plants. eLife 2: e00269.

Zhang X, Henriques R, Lin S-S, Niu Q-W, Chua N-H. 2006. Agrobacterium-mediated transformation of Arabidopsis thaliana using the floral dip method. Nature protocols 1: 641–646.

Zhang L, Hu YB, Wang H Sen, Feng SJ, Zhang YT. 2015. Involvement of miR156 in the regulation of vegetative phase change in plants. Journal of the American Society for Horticultural Science 140: 387–395.

