## Supplemental tables and figures for "Differential submergence tolerance between juvenile and adult Arabidopsis plants involves the ANAC017 transcription factor"

**Supplemental information to:**

**“Variable submergence tolerance across vegetative stages of *Arabidopsis thaliana* involves the ANAC017 transcription factor”**

Liem T. Bui^1^, Vinay Shukla^1^, Federico M. Giorgi^2^, Alice Trivellini^3^, Francesco Licausi^1,2^ and Beatrice Giuntoli^1,2^

^1^Plantlab, Institute of Life Sciences, Scuola Superiore Sant’Anna, Italy

^2^Pharmacology and Biotechnology Department, University of Bologna, Italy

^3^Biology Department, University of Pisa, Italy

**Content:**

**Figures S1-S7**

**Tables S1-S6**

**Supplemental Figures**

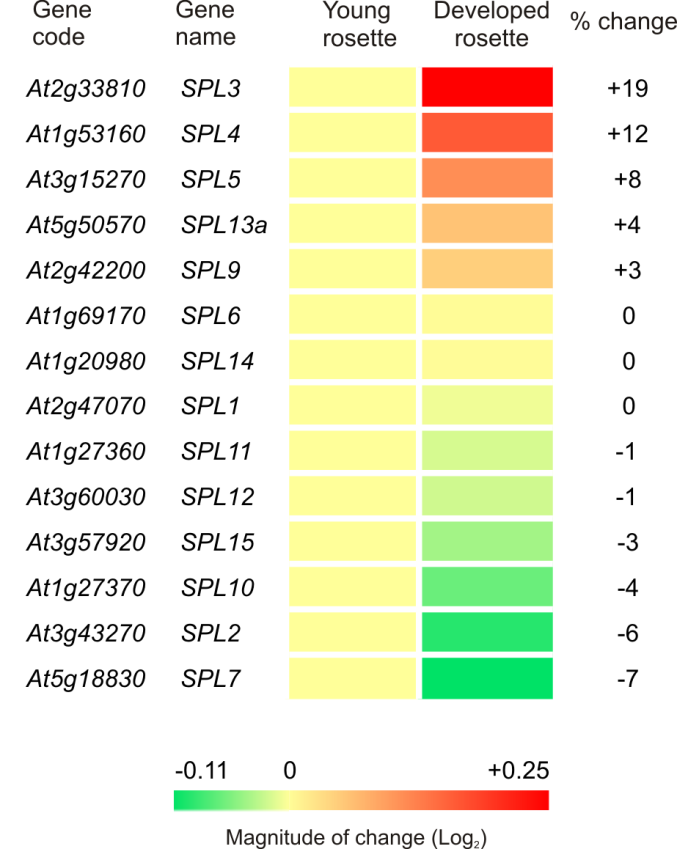

**Fig. S1 *SPL* family gene expression in juvenile or adult plant.** Expression profile at the young or developed rosette stage of all *SPL* family genes, extracted from the public microarray datasets available on the Genevestigator platform (Hruz et al., 2008). Among the gene family members, only *SPL8* was not represented on the Affymetrix probeset array. The “Development” tool built in the platform recapitulates in the “young” and “developed rosette” categories shoot samples falling, respectively, in the juvenile stage of Arabidopsis development or in the adult pre-bolting stage. Absolute expression signals were Log_2_-trasformed and made relative to the young rosette sample (Log_2_=0) for every gene. The fold change in absolute expression in developed rosettes has also been calculated, in the last column.

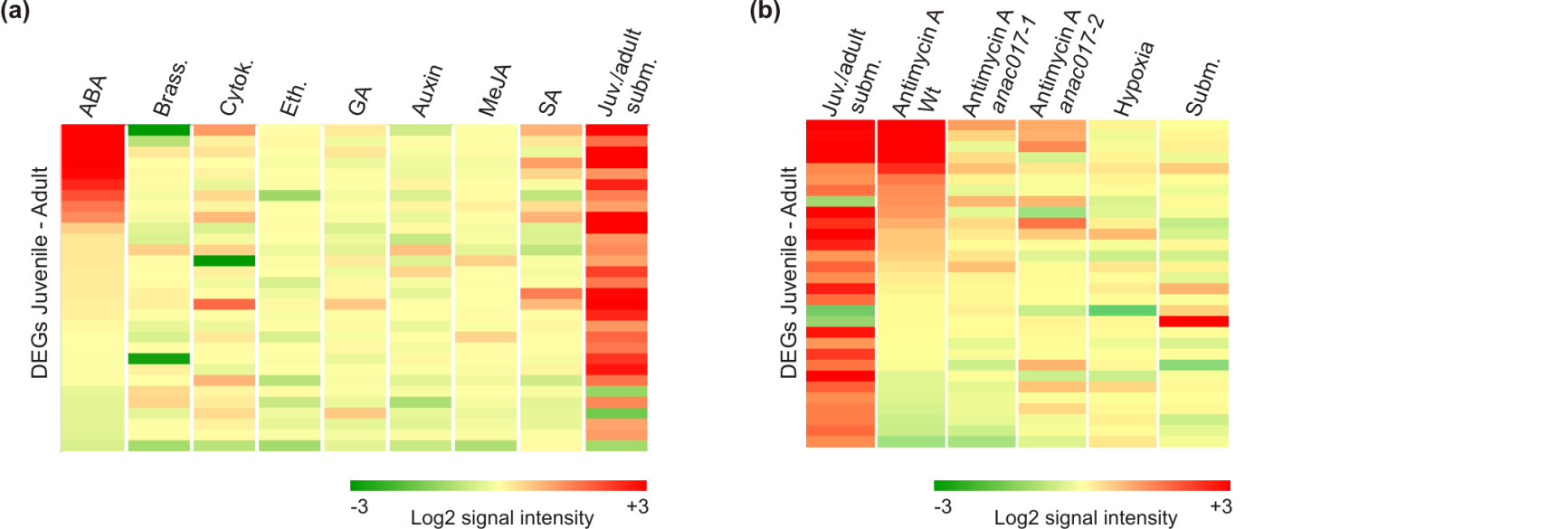

**Fig. S2. Regulation of the juvenile-specific genes in response to external perturbations.** Expression patterns of the juvenile-specific genes, identified with the global gene expression approach presented in Figure 5 and listed in Table 1, across publicly available microarray datasets related to **(a)** exogenous hormone treatments, or **(b)** treatment with antimycin A, hypoxia (1% for 1.5 h) or submergence (24 h). Data were extracted from Genevestigator. Only 30 of the 35 genes obtained from our analysis could be mapped on the Affymetrix platform. Details about the individual experiments presented here can be found in Table S6. The effect of antimycin A was evaluated in two independent *anac017* T-DNA mutants, in addition to a background genotype (“Wt”).

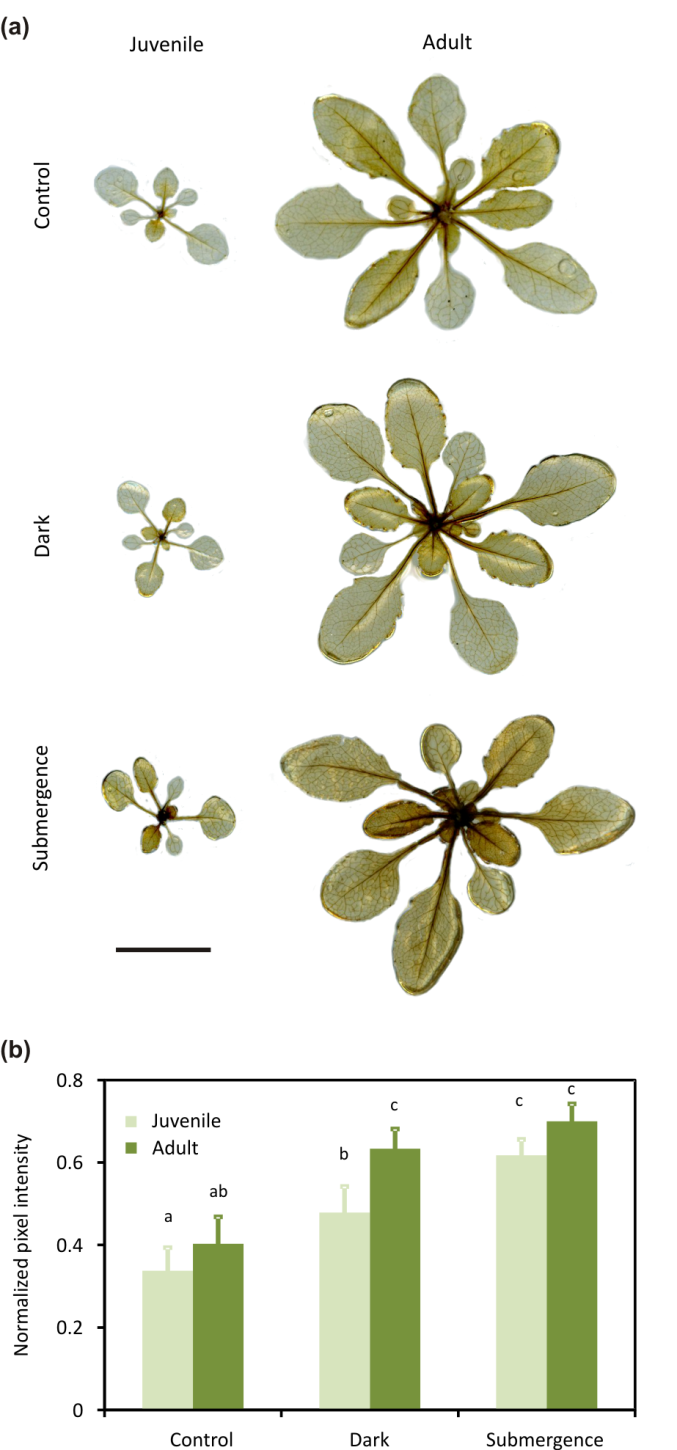

**Fig. S3. Histochemical assessment of ROS production in juvenile and adult plants.** **(a)** Representative images of DAB staining in two and three week-old plants under different stress treatments. Darker colour indicates higher presence of ROS. Scale bar, 2 cm. **(b)** Quantification of coloured pixel intensity in DAB stained plants in the different conditions. The procedure adopted is detailed in the Materials and Methods section. Data are mean ± S.D. (n=6). Different letters mark statistically significant means after two-way ANOVA (p<0.05) and Tukey post-hoc test.

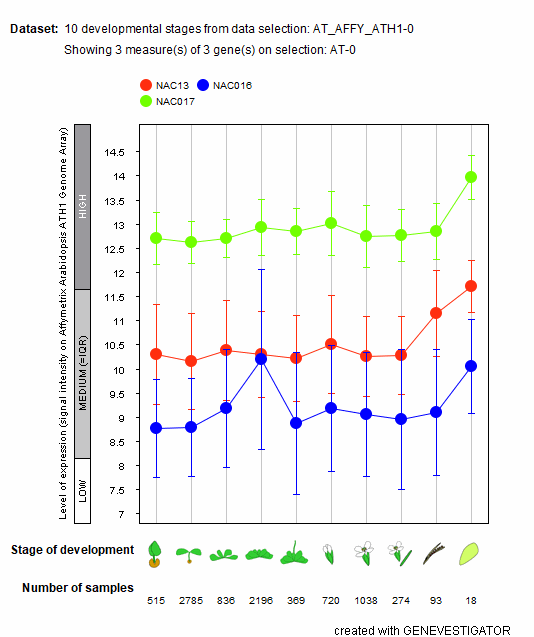

**Fig. S4 Profile of expression of *ANAC013* (red line), *16* (blue), and *17* (green) across ten stages Arabidopsis development.** Data were extracted from the “Development” platform of Genevestigator (Hruz et al., 2008), where they are presented as averaged normalized Log_2_-transformed signal intensities.

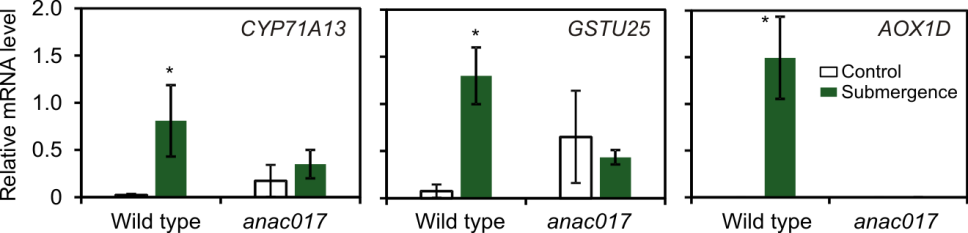

**Fig. S5 Expression of three submergence-responsive genes (*CYP71E*, *GST25* and *AOX1D*) in the wild type and the *anac017-1* mutant.** Juvenile plants were used. Submergence was protracted for 24 h. Data are mean ± S.D. (n=4). Asterisks indicate statistically significant differences between treatments in each genotype, evaluated by means of a t-test (p<0.05).

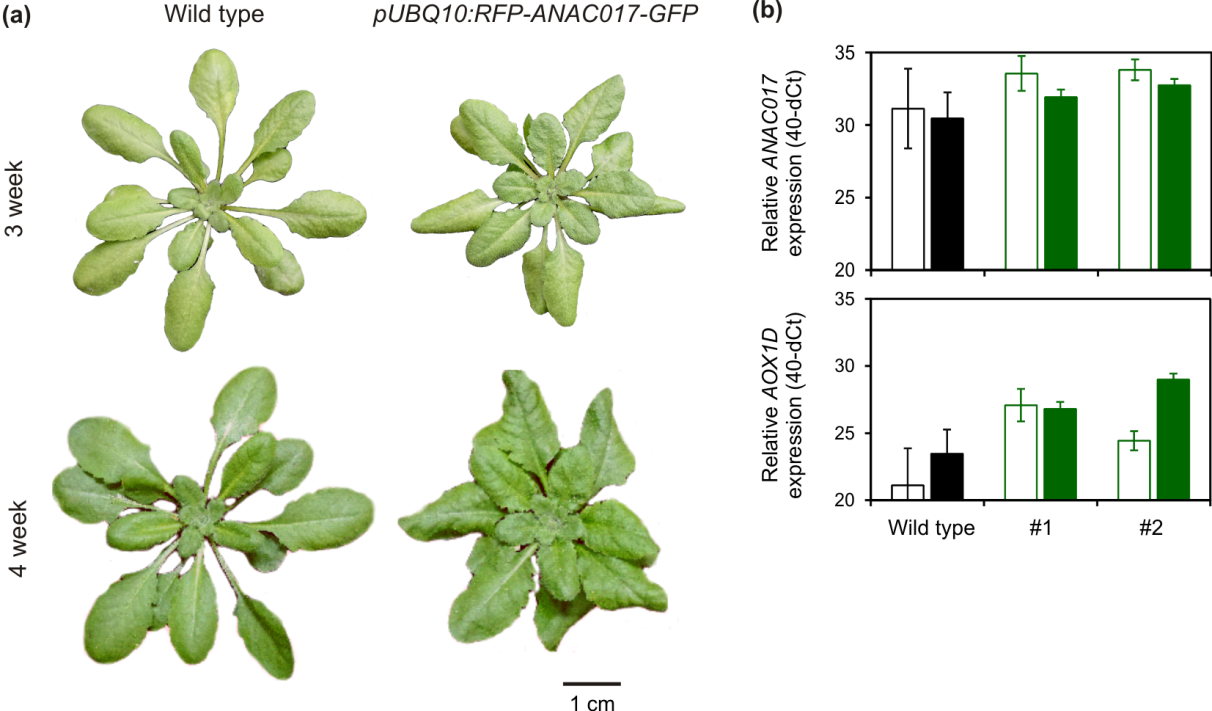

**Fig. S6 Phenotypical and molecular evaluation of *pUBQ10:RFP-ANAC017-GFP* transgenic plants*.* (a)** Vegetative phenotype of adult *pUBQ10:RFP-ANAC017-GFP* plants. Representative soil-grown individuals were depicted along Col-0 wild types at two ages of vegetative development. Scale bar, 2 cm. **(b)** *ANAC017* expression was evaluated in adult plants from two independent T_3_ lines (in green), along with the juvenile-specific target *AOX1D*, in aerobic (empty columns) or submerged shoot samples (solid columns). Data are mean ± S.D. (n=3) of 40-dCt amplification values obtained using *UBQ10* as housekeeping. Submergence was protracted for 24 h. Line #1 was kept for further experiments.

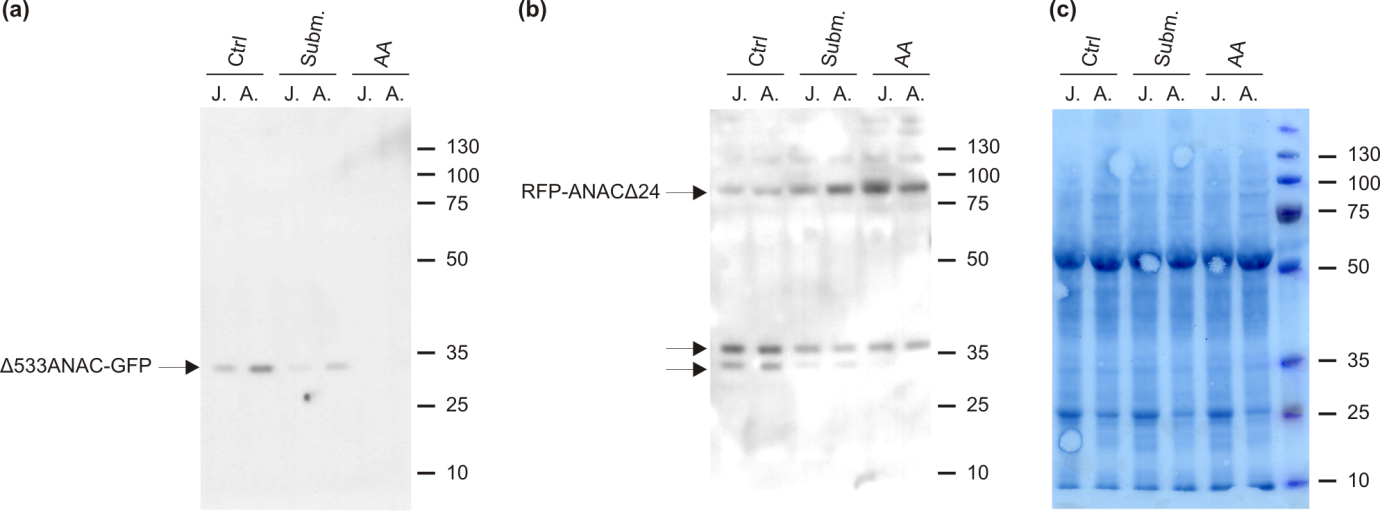

**Fig. S7 Full immunoblot images related to Fig. 8d. (a)** Membrane signal upon hybridization with an anti-GFP primary antibody. The arrow point at the hypothetical band corresponding to the C-terminal ER-tethered fragment of the ANAC017 reporter construct. **(b)** Membrane signal upon hybridization with an anti-RFP primary antibody. The top arrow indicates the hypothetical band from the N-terminal fragment putatively targeted to the nucleus. Two smaller bands, indicated by the lower arrows, might represent unspecific hybridization, as well as more minute RFP-ANAC017 peptides originated by further cleavage of the protein. **(c)** Amido black staining of the PVDF membrane. Molecular weight markers (kDa) are displayed. J., juvenile; A., adult. Ctrl, 24 h extended darkness; Subm., 24 h dark submergence; AA, 6 h treatment with 10 µM antimycin A.

**Supplemental Tables**

**Table S1. Primers used for real time qPCR measurements of gene expression.**

| **AGI code** | **Primer name** | **Primer sequence** |
| --- | --- | --- |
| AT1G01480 | ACS2_F | ATGGGTCTTGCAGAGAATCAGC |
|  | ACS2_R | ACCTTCAAGGGTGCAAATAGAAGC |
| At1g77120 | ADH_F | TATTCGATGCAAAGCTGCTGTG |
|  | ADH_R | CGAACTTCGTGTTTCTGCGGT |
| AT1G34190 | ANAC017_F | TCAGGTGTGATGCCTGATTCCAC |
|  | ANAC017_R | AGCCGATGAGAACTGGCTTGTTG |
| AT1G32350 | AOX1D_F | AGAGGCTGAGAACGAGCGTATG |
|  | AOX1D_R | GCTCGTTCGTACCATTTGGGTTG |
| AT1G07890 | APX1_SG_Fw | TGCTGATTTCCATCAGCTTG |
|  | APX1_SG_Rev | GTTGGGGCTTGTCCTCTCTT |
| AT4G34210 | ASK11_F | AAGACGATTGCGTTGCTGATGG |
|  | ASK11_R | CCATGAACTTCTCGTCCCAGTTG |
| AT1G68620 | AT1G68620_F | AGCAACATGGAGATGTGCGATGG |
|  | AT1G68620_R | GACCAACGCCTTTGTGCAACAC |
| AT3G10320 | AT3G10320_F | ATTGCGCTAAGAGATCGAAGCC |
|  | AT3G10320_R | AATCGATGAGAAACAGAGCAGAGG |
| AT1G76410 | ATL8_F | AGACGAGCTTAGGGTGTTC |
|  | ATL8_R | CGCCACATTTATGACACCTG |
| AT2G30770 | CYP71A13_F | GTGCTTCGGTTGCATCCTTCTC |
|  | CYP71A13_R | CGCCCAAGCATTGATTATCACCTC |
| AT3G47340 | DIN6_F | AAGGTGCGGACGAGATCTTTGG |
|  | DIN6_R | ACTTGTGAAGAGCCTTGATCTTGC |
| AT2G38340 | DREB19_F | TGAGTCACCGTGGTGCAAACTC |
|  | DREB19_R | CTGCAGTAGCAAACGTGCCAAG |
| AT1G17180 | GSTU25_F | TCAGCATTGAAGCCGAGTGTCC |
|  | GSTU25_R | TAGCCACACTCTCTCTCTCCACAC |
| At2g16060 | HB1_F | TTTGAGGTGGCCAAGTATGCA |
|  | HB1_R | TGATCATAAGCCTGACCCCAA |
| At3g10040 | HRA1_F | ACAACCACCGCAACAGAATCC |
|  | HRA1_R | TCTCCGCAATTCTCGCCAT |
| AT3G59940 | KMD40_F | TGGAACGATGATGGTGAAGA |
|  | KMD40_R | AACCAGAGGGAGTGTTGACG |
| AT4G33070 | PDC_F | CACAGAATCTTCAATGTTCTTACC |
|  | PDC_R | CCATGATAAAGCGTACATGGAA |
| At3g11410 | PP2CA_F | TCCTCTCTCCGTAGATCACAAGCC |
|  | PP2CA_R | ACTCCAAGAACCCTAGCTCCATCC |
| AT1G43800 | HUP7_F | TTGGCAACCCGCTTCTTTCTTACC |
|  | HUP7_R | TTTCCCTCAGCTCACGAACCTG |
| AT5G43270 | SPL2_F | AGTCGTTGTGAGTGGCGTAGAG |
|  | SPL2_R | ACTCAGAGAGACAGTGGAACCTG |
| AT2G33810 | SPL3_F | CTTAGCTGGACACAACGAGAGAAGGC |
|  | SPL3_R | GAGAAACAGACAGAGACACAGAGGA |
| AT1G53160 | SPL4_F | TTTCTCTCAGGACTTAACCAACGC |
|  | SPL4_R | CTTGGAGGTCATGAAACCTACTGC |
| AT3G15270 | SPL5_F | AAGGCATCTGCTGCGACTGTTG |
|  | SPL5_R | TCTGGTAGCTCATGAAACCTGCTG |
| AT2G42200 | SPL9_F | TGTGGCTGGTATCGAACAGAGG |
|  | SPL9_R | TTCCGGAAGCTGATGAAACCTG |
| At4g05320 | UBQ_F | GGCCTTGTATAATCCCTGATGAATAAG |
|  | UBQ_R | AAAGAGATAACAGGAACGGAAACATAGT |

**Table S2. Primers used for the MSRE-qPCR analysis of the selected genomic loci.** The methylation-sensitive restriction enzymes used for the MSRE-qPCR are indicated in column “RE”. The position of the RE relative to the translation start site is given in brackets.

| **Name of**  **target region** | **RE** | **Primer name** | **Primer sequence** |
| --- | --- | --- | --- |
| *ACS2_1* | HpaII (-1069) | AT1G01480_metF1 | 5’-GAACTAGGTCTAGTCACGTG |
|  |  | AT1G01480_metR1 | 5’-AGTCAATCACATGCTCTACGT |
| *ACS2_2* | HpaII (+11) | AT1G01480_metF2 | 5’-TGAGAATTGACACAGCAAATGG |
|  |  | AT1G01480_metR2 | 5’-AGTTCTCTCCGTGTTGATTGT |
| *AOX1D* | HpaII (+35, +63) | AT1G32350_metF | 5’-TTTACCGCACTCTTCGACCG |
|  |  | AT1G32350_metR | 5’-AGATGTCCCCTGAATCCTCCA |
| *ASK11_1* | SalI (-1444) | AT4G34210_metF1 | 5’-GCTTAGTGTGTCTTTGCAACTG |
|  |  | AT4G34210_metR1 | 5’-AACAGTACACCCATCCACGC |
| *ASK11_2* | SalI (+192) | AT4G34210_metF2 | 5’-TGGAATCCCTCTTGCAAACGT |
|  |  | AT4G34210_metR2 | 5’-TCGTCCCAGTTGTTGAGATCC |
| *CYP71A3* | HpaII (+172) | AT2G30770_metF | 5’-GAAACCTCCACCAGCTGAGC |
|  |  | AT2G30770_metR | 5’-CACGGCCAAAATGAAGGAGC |
| *DREB19A* | HpaII (-1021) | AT2G38340_metF | 5’-GCACAAATAATTCGATGAGCAGC |
|  |  | AT2G38340_metR | 5’-CCTTTGCCGAACTGGACTCG |
| *GSTU25* | HpaII (+146,+164,+188) | AT1G17180_metF | 5’-CTCCTCGAGATGAATCCGGT |
|  |  | AT1G17180_metR | 5’-AGGCCAAACTTCGTCGATGT |
| *MUCI2_1* | HpaII (-749) | AT3G10320_metF1 | 5’-TCTTGGATCATGCAGTCTTTCAC |
|  |  | AT3G10320_metR1 | 5’-TGAGGTCCAGCTCATTGTGA |
| *MUCI2_2* | HpaII (+1188) | AT3G10320_metF2 | 5’-GCTAAGAGGCCAAAACTAGCG |
|  |  | AT3G10320_metR2 | 5’-CAAACCCGATCCTCTGAGCC |
| *At1g68620* | SalI (+788) | AT1G68620_metF | 5’-GGTGGAGGATGTCTTTGCCA |
|  |  | AT1G68620_metR | 5’-CACACCAGCGTCCTTGTCA |

**Table S3. Primers used for the ChIP-qPCR assessment of histone 3 methylation on selected juvenile-specific gene loci.**

| **Name of**  **target region** | **Primer name** | **Primer sequence** |
| --- | --- | --- |
| *AOX1D_1* | At1g32350_chF1 | 5'-GCCACTTGCCCAATGTTCGG |
|  | At1g32350_chR | 5'-ACTTTACCATCGGCCCTTCGG |
| *AOX1D_2* | At1g32350_chF2 | 5'-CTAGAGACGGTGGCTGCGGT |
|  | At1g32350_chR | 5'-CCGCTATGTTCGAACCTCCGG |
| *ASK11* | At4g34210_chF | 5'-AAGACGATTGCGTTGCTCATGG |
|  | At4g34210_chR | 5'-CCATGAACTTCTCGTCCCAGTTG |
| *ACS2_1* | At1g01480_chF1 | 5'-GGAGCCACCGGAGCCAATGA |
|  | At1g01480_chR | 5'-GCGGCATAGTACGGGGAGGG |
| *ACS2_2* | At1g01480_chF2 | 5'-AGCTCAATGTGTCTCCTGGCTCT |
|  | At1g01480_chR | 5'-ATCCGTCCAAGCGCCACATG |
| *CYP71A13_1* | At2g30770_chF1 | 5'-CGTTCCCTCCGGTCCCTCAG |
|  | At2g30770_chR | 5'-CCTCTTGAGCTGCTTCACCGG |
| *CYP71A13_2* | At2g30770_chF2 | 5'-CGAGTAGAGTTGCGTTGGGAAGA |
|  | At2g30770_chR | 5'-TCCATGATCTGCCTCACTCGCT |
| *DREB19* | At2g38340_chF | 5'-TGAGCCAGTGGAAGCGACGT |
|  | At2g38340_chR | 5'-TCGAACCTTTGGCTTGAACCCT |
| *GSTU25* | At1g17180_chF | 5'-TCTGGCCGAGCATGTTTGGA |
|  | At1g17180_chR | 5'-TCCGGCTTTTGTTCCACAGATCT |
| *MUCI2_1* | At3g10320_chF1 | 5'-TTGCGCTAAGAGATCGAAGCCT |
|  | At3g10320_chR | 5'-ACAGAGCAGAGGGGTAAGGGA |
| *MUCI2_2* | At3g10320_chF2 | 5'-ACCGTCGATCCTTCTCAGATGCA |
|  | At3g10320_chR | 5'-GCCTGTCCAGTCTGTTGATTCGT |
| *At1G68620_1* | At1g68620_chF1 | 5'-GGCGTGACTTGCTCCGATGT |
|  | At1g68620_chR | 5'-ACTTGGTCGTGGTCATCGGGA |
| *At1G68620_2* | At1g68620_chF2 | 5'-GTCCGACGCATGGTGGAGCA |
|  | At1g68620_chR | 5'-AGCGTCCTTGTCACCGTCGA |

**Table S4. Numeric expression values of marker genes for submergence-related processes in treated juvenile (“J.”) or adult plants (“A.”).** For each gene, data were normalized on the expression value of juvenile plants before the treatment (T_0_). Values are mean ± S.D. (n=4). Asterisks indicate statistically significant differences between juvenile and adult plant samples at each time point of treatment, evaluated by means of independent t-tests (p<0.05). The same data are displayed as a heatmap in Fig. 4.

| **Gene name** | **Age** | **T_0_** | **12 h subm.** | **12 h subm + reox.** | **24 h subm.** | **24 h subm + reox.** |
| --- | --- | --- | --- | --- | --- | --- |
| *ADH* | Juvenile | 1.00±0.16 | 235.50±213.41 | 7.17±1.81* | 717.75±350.24 | 11.94±3.77 |
|  | Adult | 1.39±0.79 | 157.25±162.80 | 12.15±1.92 | 318.50±49.02 | 19.51±6.16 |
| *PDC* | Juvenile | 1.00±0.13 | 2066.84±2073.89 | 8.54±6.93 | 3164.27±858.02 | 31.55±17.75 |
|  | Adult | 0.90±0.37 | 1001.26±822.68 | 6.04±4.86 | 2760.83±1102.40 | 41.98±27.29 |
| *HB1* | Juvenile | 1.00±0.12 | 1807.11±949.29 | 5.94±1.60 | 2952.06±482.31 | 26.62±16.54 |
|  | Adult | 1.53±0.88 | 1592.02±396.69 | 5.04±2.12 | 2358.03±781.02 | 50.1518.61 |
| *HUP7* | Juvenile | 1.00±0.44 | 4444.78±4131.10 | 1.51±1.24 | 2177.54±286.79 | 16.59±11.16 |
|  | Adult | 2.09±1.05 | 2501.88±2326.14 | 3.51±1.80 | 2905.94±492.78 | 30.39±5.18 |
| *HRA1* | Juvenile | 1.00±0.33 | 39.12±37.80 | 0.512±0.16 | 5.58±0.75 | 0.41±0.10 |
|  | Adult | 0.52±0.14 | 35.97±21.08 | 0.59±0.24 | 17.87±10.09 | 0.48±0.04 |
| *APX* | Juvenile | 1.00±0.61 | 0.48±0.07 | 1.07±0.75 | 28.14±4.51* | 44.72±26.34 |
|  | Adult | 0.63±0.79 | 0.31±0.20 | 0.92±0.34 | 11.74±6.58 | 35.67±15.84 |
| *DIN6* | Juvenile | 1.00±0.45 | 6.19±2.59 | 1.50±1.00 | 35.98±10.69* | 23.13±4.53 |
|  | Adult | 0.84±0.79 | 7.27±1.17 | 5.14±2.96 | 154.10±55.19 | 23.71±14.97 |
| *At1g76410* | Juvenile | 1.00±0.28 | 0.30±0.08 | 0.55±0.13 | 0.52±0.15 | 0.74±0.14 |
|  | Adult | 0.48±0.39 | 0.34±0.07 | 0.51±0.051 | 1.53±0.68 | 1.1±0.08 |
| *TPS8* | Juvenile | 1.00±0.29* | 1.59±0.42 | 0.58±0.10 | 0.79±0.14* | 0.94±0.10 |
|  | Adult | 0.35±0.14 | 1.51±0.32 | 0.80±0.21 | 1.80±0.40 | 0.92±0.21 |
| *KMD4* | Juvenile | 1.00±0.28* | 0.39±0.06 | 0.84±0.19 | 0.14±0.02* | 1.42±0.29 |
|  | Adult | 0.38±0.08 | 0.42±0.06 | 0.96±0.30 | 0.31±0.10 | 1.40±0.29 |

**Table S5 (separate file). Global gene expression analysis in shoot samples from juvenile and adult Col-0 plants after 24 h dark submergence. (a)** Full data set. **(b)** Filtered data set showing genes differentially regulated by the treatment (Log_2_FC threshold=2) only in juvenile plants (1727 probe sets), or **(c)** only in adult plants (1595 probe sets). **(d)** Stringent data set selection of genes differentially expressed (Log_2_FC threshold=1)) between juvenile and adult plants under submergence with adj. p (Sub2/Sub3)<0.05 (35 probe sets). This selection is also reported in Table 1.

**Table S6 (separate file). Expression profile of juvenile-specific genes in public microarray experiments analysed with the aid of Genevestigator (Hruz. et al. 2008). (a)** Hormone-related experiments. **(b)** Antimycin A and low oxygen experiments. Log_2_FC values are reported.
