## Supplementary material for "Differential submergence tolerance between juvenile and adult Arabidopsis plants involves the ANAC017 transcription factor": Table 1

**Table 1. Juvenile-specific genes during submergence identified from the global transcript analysis.**

| **AGI code** | **Annotation** | **Log_2_FC (Sub2/Sub3)** |
| --- | --- | --- |
| *AT5G55150* | Protein of unknown function (DUF295) | 4.36 |
| *AT1G17180* | GSTU25, glutathione S-transferase TAU 25 | 3.98 |
| *AT1G26390* | FAD-binding Berberine family protein | 3.43 |
| *AT5G59330* | Bifunctional inhibitor/lipid-transfer protein/seed storage 2S albumin superfamily protein | 3.40 |
| *AT2G38240* | 2-oxoglutarate (2OG) and Fe(II)-dependent oxygenase superfamily protein | 3.39 |
| *AT1G32350* | AOX1D, alternative oxidase 1D | 3.38 |
| *AT3G29250* | SDR4, NAD(P)-binding Rossmann-fold superfamily protein | 2.99 |
| *AT2G04070* | MATE efflux family protein | 2.93 |
| *AT1G55010* | PDF1.5, plant defensin 1.5 | 2.89 |
| *AT5G40010* | AATP1, AAA-ATPase 1 | 2.76 |
| *AT3G10320* | Glycosyltransferase family 61 protein | 2.65 |
| *AT2G36440* | Unknown protein | 2.59 |
| *AT4G22520* | Bifunctional inhibitor/lipid-transfer protein/seed storage 2S albumin superfamily protein | 2.51 |
| *AT1G70850* | MLP34, MLP-like protein 34 | 2.40 |
| *AT5G46960* | Plant invertase/pectin methylesterase inhibitor superfamily protein | 2.27 |
| *AT3G09260* | BGLU23, glycosyl hydrolase superfamily protein | 1.80 |
| *AT2G35730* | Heavy metal transport/detoxification superfamily protein | 1.71 |
| *AT3G53190* | Pectin lyase-like superfamily protein | 1.64 |
| *AT2G23260* | UGT84B1, UDP-glucosyl transferase 84B1 | 1.59 |
| *AT2G47010* | Unknown protein | 1.58 |
| *AT1G51140* | Basic helix-loop-helix (bHLH) DNA-binding superfamily protein | 1.50 |
| *AT3G56891* | Heavy metal transport/detoxification superfamily protein | 1.42 |
| *AT4G34160* | CYCD3;1, CYCLIN D3;1 | 1.42 |
| *AT4G02290* | GH9B13, glycosyl hydrolase 9B13 | 1.38 |
| *AT1G01480* | ACS2, 1-amino-cyclopropane-1-carboxylate synthase 2 | 1.27 |
| *AT1G15630* | Unknown protein | 1.23 |
| *AT1G80080* | AtRLP17TMM, leucine-rich repeat (LRR) family protein | 1.22 |
| *AT2G39690* | Protein of unknown function (DUF547) | 1.20 |
| *AT1G07880* | ATMPK13, protein kinase superfamily protein | 1.18 |
| *AT1G61120* | TPS4, terpene synthase 4 | 1.13 |
| *AT1G07180* | ATNDI1, alternative NAD(P)H dehydrogenase 1 | 1.06 |
| *AT2G32010* | CVL1, CVP2 like 1 | 1.06 |
| *AT5G49360* | BXL1, beta-xylosidase 1 | -1.08 |
| *AT5G25180* | CYP71B14, cytochrome P450, family 71, subfamily B, polypeptide 14 | -1.22 |
| *AT1G50040* | Protein of unknown function (DUF1005) | -1.63 |
